# GxE and changes to the genetic architecture of flowering time in an overwintering environment

**DOI:** 10.64898/2026.10.30.762947

**Authors:** Amanda L. Peake, John R. Stinchcombe

## Abstract

Characterizing how the expression of genetic variation changes across environments is essential for understanding evolution in complex environments, the evolution of phenotypic plasticity, and the genetic architecture of adaptation. We investigated how the phenotypic distribution, large-effect loci, and polygenetic architecture of flowering time in *Arabidopsis thaliana* changes when grown in an ecologically realistic environment compared to standard chamber conditions. We grew 479 *A. thaliana* accessions in an overwintering common garden that have previously been phenotyped in growth chambers by the 1001 Genomes Project (2016). Our findings suggest there is extensive GxE for flowering time between the overwintering common garden and the growth chamber environments. None of the large effect loci that we identified using a multivariate Genome Wide Association Study (GWAS) were shared across growth conditions. We find evidence of allelic differential sensitivity, conditional neutrality, and antagonistic pleiotropy contributing to GxE for flowering time between growth conditions. Our results also suggest that the effects of genetic variation across multiple flowering time regulatory pathways is revealed when grown in a more ecologically realistic environment. The effect sizes for the most significant 1% of SNPs are moderately correlated between the outdoor and growth chamber conditions which indicates a partially shared genetic basis. Overall, our findings highlight how making inferences about the genetic basis and evolution of complex traits may be dependent on conducting experiments in ecologically realistic environments.

**Article Summary:** A crucial aspect of characterising the genetic basis of complex traits is knowing how genetic variation is expressed differently across environmental conditions. This research compares the genetic architecture of *Arabidopsis thaliana* flowering time between an ecologically realistic environment to standard lab conditions. Although the genetic basis of flowering time is partially shared between environments, large-effect loci differ in both magnitude and direction of effects across environments. These findings highlight how phenotyping traits in realistic environments is essential for understanding the contribution of individual loci, roles of different genetic pathways, and evolutionary consequences of phenotypic variation in complex traits.

## Introduction

A fundamental goal in evolutionary genetics is to characterize the genetic basis of ecologically important traits (Conner and Hartl 2004). Understanding the genetic basis of these traits, however, is challenging, because the phenotypic variation observed in nature is due to a combination of genotype, environment, and genotype-by-environment interactions (Lynch and Walsh 1998). In addition, organisms frequently respond to a combination of environmental cues in nature (temperature, daylength, moisture or humidity, light availability, etc), which suggests that components of phenotypic and genetic variation might only be revealed when we measure phenotypes in ecologically realistic environments (El-Soda et al. 2014; Wood and Brodie 2015). Genetic variants associated with a trait in one environment can have no influence, a reduced impact, or even an opposite effect in a different environment due to conditional neutrality, allelic differential sensitivity, and environmental antagonistic pleiotropy (Des Marais et al. 2013). Resolving how the expression of genetic variation differs across environments is necessary to address fundamental questions about genetic architecture, the evolution of phenotypic traits and phenotypic plasticity, and the forces affecting the maintenance of phenotypic and genetic variation (Lynch and Walsh 1998). Here, we investigate how the genetic basis of flowering time in *Arabidopsis thaliana* changes in an outdoor overwintering common garden, and we show that the effects of genetic variation across multiple regulatory pathways are revealed when individuals are grown in a more ecologically realistic environment.

Flowering time is one of the most commonly studied plant traits, and rightly so, given its direct impact on plant reproductive success (Austen et al. 2017). In annual plants, like *A. thaliana* (hereafter, *Arabidopsis*), flowering often needs to be initiated within a narrow optimal window that allows individuals to have sufficient resources to successfully complete their life cycle while avoiding damage due to environmental pressures like drought and cold (Exposito-Alonso 2020). *Arabidopsis* has an extensive geographic range, suggesting that throughout its evolutionary history it has adapted to or at least experienced a wide variety of ecological and environmental conditions (1001 Genomes Consortium 2016; Lee et al. 2017). Flowering time in *Arabidopsis* can be initiated by multiple regulatory pathways, some of which respond to external stimuli such as the vernalization, temperature, and photoperiod pathways (Bouché et al. 2016); the cues for these pathways also vary along geographic and environmental gradients. Understanding how individual genetic components come together to produce phenotypic variation in flowering time requires phenotyping individuals in the presence of external stimuli that are often hard to replicate in controlled lab conditions.

Previous work in *Arabidopsis* has shown that flowering time phenotypic distributions, large-effect loci, and underlying genetic interactions change in different growth conditions. For example, Swedish accessions grown in growth chambers had different responses where some accessions flowered early and others flowered late in a 16°C growth chamber compared to a 10°C growth chamber (Sasaki et al. 2015). Similarly, changing temperature regimes in growth chambers to mimic daily fluctuating temperatures leads to a reduced influence of large effect size loci on flowering time (Burghardt et al. 2016). Recombinant-inbred lines (RILs) grown in growth chambers simulating different seasons have also revealed the presence of condition-specific epistatic interactions (Li et al. 2006). Moreover, accessions and RILs grown in the field lead to the discovery of different QTL associated with flowering time (Brachi et al. 2010). Mutants for large-effect flowering time loci also have varied effects on flowering time across multiple field sites, but more importantly the fitness effects of the flowering time mutations also differ between environments (Taylor et al. 2019). When phenotyping a large global collection of accessions in different growth chamber environments set to 10°C and 16°C some GWAS hits for flowering time were not shared between environments (1001 Genomes Consortium 2016). There is also evidence that loci under selection from a representative global sample of *Arabidopsis* accessions have different effects on fitness across field conditions with different rainfall treatments (Exposito-Alonso et al. 2019). Taken together, these results suggest that there is GxE for flowering time in *Arabidopsis* likely due to changes in the underlying genetic architecture, which has implications for plant fitness. However, most studies investigating how the genetic basis of flowering time differs between environments use RILs or mutant lines, limiting genetic resolution; a smaller subset of naturally occurring accessions, which may exclude naturally occurring genetic variation; or growth chamber environments which do not mimic natural conditions. Therefore, to better understand the genetic architecture underlying variation for *Arabidopsis* flowering time, GWAS style-studies, using a large number of genomically well-characterized and previously phenotyped samples, are needed in ecologically relevant environments.

We investigate how the phenotypic distribution, effects of individual loci, and the overall polygenic architecture for flowering time change when grown in an overwintering common garden compared to controlled lab conditions. We grew 479 accessions in a common garden that had previously been collected, sequenced, and phenotyped in growth chambers kept at a constant 10°C and 16°C (1001 Genomes Consortium 2016). We observed differences in the flowering time phenotypic distribution and mean between the overwintering common garden and growth chambers environments. None of the loci we identified as being significantly associated with flowering time were shared across all three growth conditions. We identified loci with SNP-by-environment interactions (SNPxE) due to conditional neutrality, allelic differential sensitivity, and environmental antagonistic pleiotropy. Additionally, the overall polygenetic architecture of flowering time differed between environments.

## Methods

### Accession Selection

We chose 500 accessions from the 1001 Genomes Project (Fig 1). We excluded samples collected from North America because many North American accessions within the 1001 Genomes Project collection are genetically identical owing to recent colonization by a single lineage (1001 Genomes Consortium 2016). Since we were interested in comparing differences in the genetic basis of traits between different environments, we chose 500 accessions that had relatively complete phenotypic data including flowering time from growth chambers kept at a constant 10°C and 16°C (1001 Genomes Consortium 2016).

**Figure 1.**
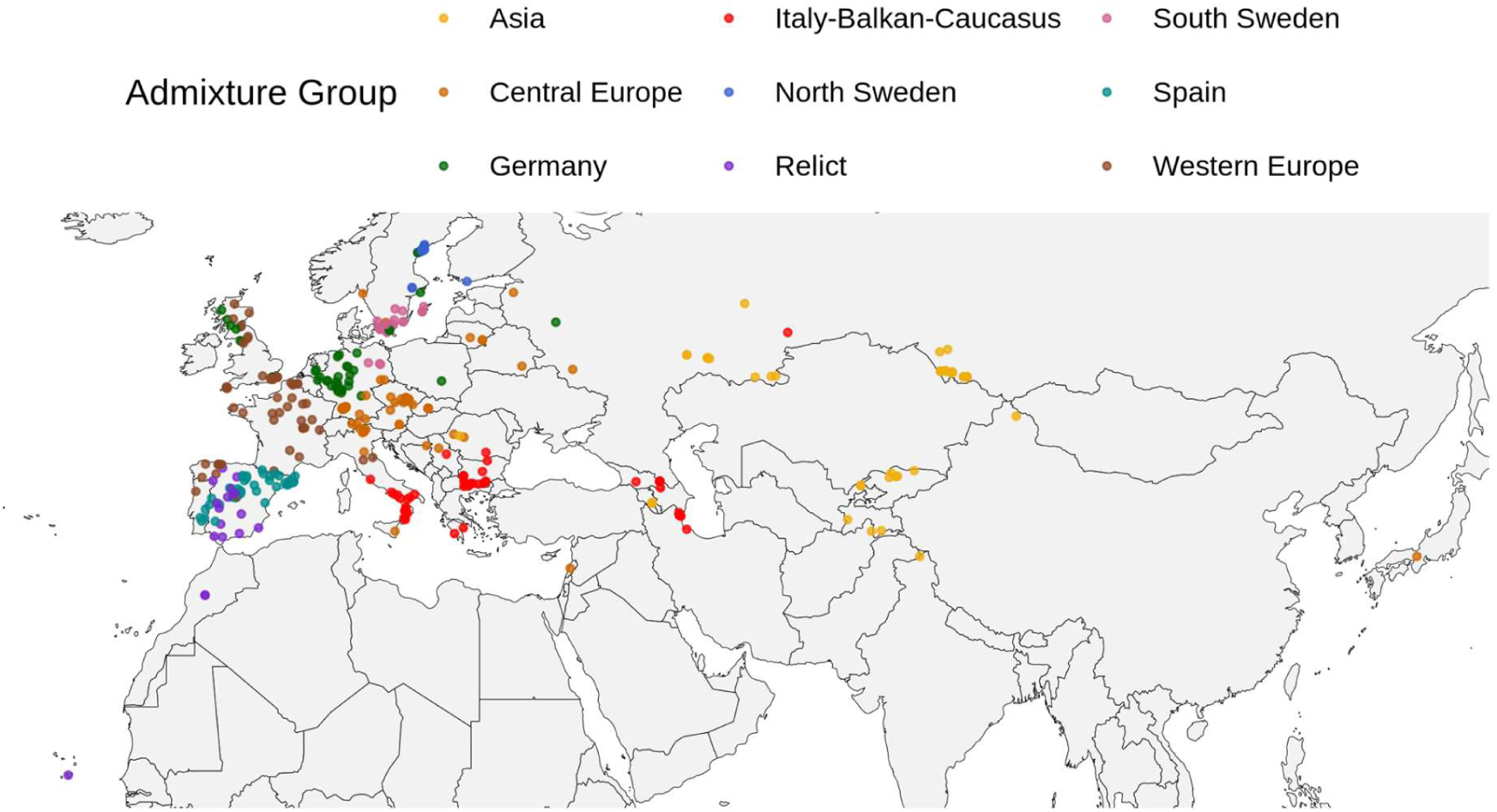
Map of sampling locations for 479 accessions colored by admixture groups as assigned by the 1001 Genomes Consortium (2016).

### Outdoor common garden

We transplanted germinates into six replicate blocks in beds filled with Promix soil on an outdoor rooftop garden of the University of Toronto Earth Sciences Building (25 Willcocks Street, Toronto, ON M5S 3B2, Canada). If there were more than six replicates available for transplanting for a given accession, we prioritized individuals that were more established. Transplanting took place over two days (October 24-25, 2022). On November 3rd 2022, we replaced any individuals (if a replacement was available) that had died due to transplant shock or had prematurely started bolting whilst in the greenhouse. We used iButtons to record surface and soil temperature in the raised beds (see Supplementary Methods).

### Flowering time

Each individual had a plant tag with a barcode indicating its stock ID number and replicate (e.g. CS76649_1). During flowering assays, we scanned all individuals that had at least one fully opened flower that day using a commercially available Tera Scanner barcode scanner. Flowering time was calculated as the elapsed days between sowing and date of the first flowering assay that an individual was recorded as flowering. Through the fall to early spring, flowering assays were conducted on a weekly basis, as long as beds were not covered by snow; the raised beds were covered with snow from approximately December 10 until after the last snowfall of the winter season on March 29. From April 12th 2023 onwards, flowering assays were conducted every other day until the end of the experiment. If weather or air quality conditions (e.g., due to forest fire smoke) did not allow for phenotyping, we conducted a flowering assay on the next possible day. We then calculated the accession mean for flowering time across all blocks. We excluded any accessions that only had a single replicate. For all downstream analyses we used the accession mean which was calculated by taking the mean across all blocks. Our final dataset consisted of 479 accessions.

### 1001 Genomes data

We downloaded flowering time phenotype data collected from growth chambers kept at 10°C and 16°C (1001 Genomes Consortium 2016) and the imputed SNP matrix (https://1001genomes.org/data/GMI-MPI/releases/v3.1/SNP_matrix_imputed_hdf5/1001_SNP_MATRIX.tar.gz) for the full 1135 accessions from the (1001 Genomes Consortium 2016). We used the python script (https://1001genomes.org/data/GMI-MPI/releases/v3.1/SNP_matrix_imputed_hdf5/h5m2csv.py) provided by the 1001 Genomes Consortium (2016) to convert the imputed SNP matrix from an h5 file into a csv file using python v3.14.2 and hdf5-mpi v1.14.6. We then used custom bash scripts to subset the SNP matrix to only include the 479 accessions included in the analysis and generate GEMMA bimbam formatted genotype, phenotype, and annotation input files. For the GEMMA genotype input file, genotypes were encoded additively (0 = homozygote reference, 1 = heterozygote, and 2 = homozygote alternative). We also downloaded the gff file for the TAIR10 reference assembly from NCBI (https://ftp.ncbi.nlm.nih.gov/genomes/all/GCA/000/001/735/GCA_000001735.2_TAIR10.1/GCA_000001735.2_TAIR10.1_genomic.gff.gz).

### Testing for GxE

Only mean flowering times for accessions from the growth chambers was available from the 1001 genomes consortium. The absence of the underlying replicate-level phenotypic data precludes the traditional two way anova testing for the effects of growth environment, genotype, and the genotype-by-environment interactions. However, we implement several related analyses to test for GxE. First, we used a Type III ANOVA with the replicate-level data from the outdoor common garden to determine if flowering time is distinguishable between accessions in the outdoor common garden. We included block as a fixed effect and accession as a random effect in the anova linear model (flowering time ∼ block + (1|accession)). We used a likelihood ratio test to determine the significance of the accession as a random effect and halved the p-value to correct for testing a random effect (Kuznetsova et al. 2015). Second, to explore the prevalence of rank changes and changes in the expression of variation across the three growth conditions, we plotted reaction norm slopes and calculated the Spearman’s rank correlations for flowering time between all three environments. We calculated the Spearman’s rank correlations in R using cor.test() function and since the data had tied ranks the p-values were calculated by specifying “exact = FALSE”. To explore whether the patterns of GxE that we observed between admixture groups were due to differences in rank order or changes in variance, we calculated the Spearman’s rank correlation within each admixture group separately. We bootstrapped the correlations with 10,000 iterations to estimate bias-corrected and accelerated 95% confidence intervals. We corrected rank correlation p-values to account of multiple pairwise comparisons between environments using a Bonferroni correction (*p_adj_* = *p_raw_*\*3) and all adjusted p-values were capped at a maximum of one. Third, we used a paired t-test between accession means in each growth condition and adjusted p-values for multiple pairwise comparisons (*p_adj_* = *p_raw_*\*3). These analyses test the null hypothesis that phenotypic plasticity (the difference in mean flowering time between conditions) across accessions is on average zero. We note that non-zero plasticity can exist without GxE (i.e., in the presence of equal environmental effects across all genotypes), but that the absence of significant phenotypic plasticity in the presence of GxE is a much more restrictive scenario.

### Multivariate GWAS

We conducted a multivariate GWAS using GEMMA v0.98.5 (Zhou and Stephens 2012; Zhou and Stephens 2014) to determine if there is a shared genetic basis for flowering time between the growth chambers and the outdoor common garden experiment. We included flowering time across all three environments (10°C growth chamber, 16°C growth chamber, and outdoor common garden) as a matrix of response variables in the multivariate GWAS. We controlled for population structure by including a centered relatedness matrix as a random effect. The centered relatedness matrix was generated using GEMMA with the same SNP set used in the GWAS. The GWAS included 1,833,927 SNPs after filtering for a minor allele frequency of 0.05. Since the multivariate GEMMA model does not generate marginal p-values for all fixed effects, we conducted a univariate GWAS for all three environments to see which environment was driving the peak in the multivariate GWAS. The univariate GWAS models were run using the same relatedness matrix and minor allele frequency filter as the multivariate model. We used likelihood ratio tests to calculate test statistics for both the multivariate and univariate GWAS linear mixed models. We corrected for multiple comparison testing with both a Bonferroni and false-discovery rate (FDR) of 5% correction.

To identify candidate genes within a GWAS peak region, we grouped SNPs that surpassed the Bonferroni corrected threshold in any of the LMM GWAS into peaks if they fell within 20 Kb of each other. We downloaded the TAIR10 annotation gff file from https://www.arabidopsis.org/api/download-files/download?filePath=Maps/gbrowse_data/TAIR10/TAIR10_GFF3_genes.gff. We then identified candidate genes that fell within 10 kb of each GWAS peak. We limited the list of candidate genes to those that have previously been experimentally validated by some form of genetic manipulation in *Arabidopsis*.

To investigate if effect sizes differed between environments, we tested whether effect sizes for the top 1% of SNPs in the multivariate GWAS were correlated across environments. We expected that effect sizes would be highly correlated across two environments if flowering time had a similar genetic basis in both environments. The multivariate output from GEMMA does not include standard error for effect sizes. Therefore, we identified the top 1% of most significant SNPs from the multivariate GWAS but then used the effect sizes and standard errors from the univariate GWAS for comparisons between environments. Since GWAS are known to be biased towards overestimating the effect sizes of significant SNPs (Göring et al. 2001), we also calculated the Spearman’s rank correlation that should be more robust to the magnitude of individual effect sizes (e.g., overestimates of significant SNP effects, or under-estimates of non-significant SNP effects). We calculated both the Pearson’s and Spearman’s rank correlations. Given the presence of values with tied ranks, p-values for the Spearman’s rank correlation were calculated by specifying “exact = FALSE”. We corrected correlation p-values to account of multiple pairwise comparisons between environments using a Bonferroni correction (*p_adj_* = *p_raw_*\*3).

Next, we evaluated how large effect loci were contributing to GxE. We determined if significant SNPs from the multivariate GWAS had different effect sizes across the different environments (i.e. evidence of SNPxE) and then categorised the nature of SNPxE as conditional neutrality, allelic differential sensitivity, or antagonistic pleiotropy. We first calculated 95% confidence intervals for SNP effect sizes as ±1.96*SE to perform pairwise comparisons across environments for each SNP. We identified SNPs with SNPxE effects if the effect size confidence intervals in both environments did not overlap. From the SNPs with SNPxE, we identified SNPs as being conditionally neutral if the 95% confidence intervals overlapped with zero in one of the environments but not the other. Of the remaining SNPs with SNPxE, we then considered SNPs with effects in opposite directions as being antagonistic pleiotropic and SNPs with effects in the same direction but with different magnitudes to have allelic differential sensitivity.

### BSLMM

We used a Bayesian-Sparse Linear Mixed Model (BSLMM) to investigate the polygenic architecture of flowering time in each environment using the GEMMA “-bslmm 1” model. We used 500,000 burn-in steps and 5,000,000 posterior sampling steps based on trace plots of posterior distributions of hyper parameters (PVE, PGE, and n_ɣ_; Fig S4-6). We subsampled every 10th iteration of the posterior sampling steps to determine the posterior distribution for the proportion of phenotypic variation explained by the phenotype (PVE), the proportion of the genetic variance explained by major effect loci (PGE), and the estimated number of SNPs that have a non-zero effect on the trait (n_ɣ_).

### Evaluating the robustness of GWAS inferences

Flowering time in the outdoor common garden was left-skewed due to some individuals flowering before winter (Fig S7a). We took several steps to investigate how the phenotypic distribution of flowering time affected our results. First we determined if flowering time in the outdoor common garden violated the assumption of normally distributed residuals. We calculated the difference between the observed phenotype and a BSLMM predicted phenotype for each accession as a proxy for genome-wide residuals. To do so, we duplicated the genotype matrix to have a “training” set with the original outdoor common garden flowering time values and a “prediction” set with phenotypes left blank to be predicted by BSLMM. We generated a centered relatedness matrix using GEMMA with a minor allele frequency of 0.05. We then ran BSLMM for the observed data and used the output as parameters for phenotype prediction. We then calculated the difference between the observed and predicted phenotype values and evaluated normality using a shapiro wilk test and a QQ plot.

We took three approaches to test how the left-skewed distribution might be affecting our results. First, we removed individuals that strongly deviated from the QQ reference line in the QQ plot generated from the genome-wide residuals. We only retained individuals (n = 417) within -1.25 and 2 of the normalized quantile distribution shown in Fig S7d. We then reran the three LMM univariate GWAS and BSLMM as described above with the subset dataset for all three growth conditions. Second, we implemented permutation-based GWAS and third, we explored a suite of data transformations to normalize flowering time prior to analysis; we present the methods, results, and discussion of these analyses in the Supplemental Note 1.

## Results

### GxE for flowering time

To determine how genotype, environment, and gene-by-environment interactions (GxE) might be affecting flowering time phenotypic variation, we compared flowering time in the outdoor overwintering common garden to flowering time previously collected for the same accessions in growth chambers kept at a constant 10°C and 16°C from the 1001 Genomes Consortium (2016). Since replicate-level data was only available for the outdoor common garden, we first tested if there was a main effect for accession within the outdoor common garden. We found a significant effect for both accession and block within the outdoor common garden (Type III ANOVA: F_block_(5, 1980.3) = 2.46, *p*_block_ = 0.03; Likelihood ratio test: *χ*²_accession_(1) = 1216.1, adjusted *p*_accession_ < 0.001) which indicates that flowering time differs between genotypes in the outdoor common garden. We observed a greater range in the outdoor common garden experiment (accession mean flowering time from 40 to 223 days) compared to the growth chambers (accession mean flowering time from 51.25 to 149.25 days in 10°C and from 32 to 145 days in 16°C). However, the larger range in the outdoor common garden experiment is likely due to some individuals flowering before being covered by snow for winter; the majority of individuals flowered after over-wintering within a short period of time following a spike in temperature mid April (Fig S1). Collectively, these observations indicate that flowering time differs between genotypes when grown in the common garden.

We observed greater effects of the growth environment when comparing the outdoor common garden experiment and the growth chambers than just the two growth chamber conditions to each other. Although we observed significant differences in the flowering time means between all three environments, the difference between the two growth chambers was smaller than the differences between the outdoor common garden and the growth chambers (paired t-test: mean difference_outdoor, 10°C_ = 106.05, t_outdoor, 10°C_ (478) = 71.81, adjusted *p*_outdoor, 10°C_ < 0.001; mean difference_outdoor, 16°C_ = 115.35, t_outdoor, 16°C_(478) = 77.18, adjusted *p*_outdoor, 16°C_ < 0.001; mean difference_10°C, 16°C_ = 9.30, t_10°C, 16°C_(478) =14.53, adjusted *p*_10°C, 16°C_ < 0.001). We observed gradual reaction norm slopes between the 10°C and 16°C growth chambers compared to the steep reaction norm slopes between the growth chambers and the outdoor common garden (Fig 2). These results indicate that flowering time differs between different environments.

**Figure 2.**
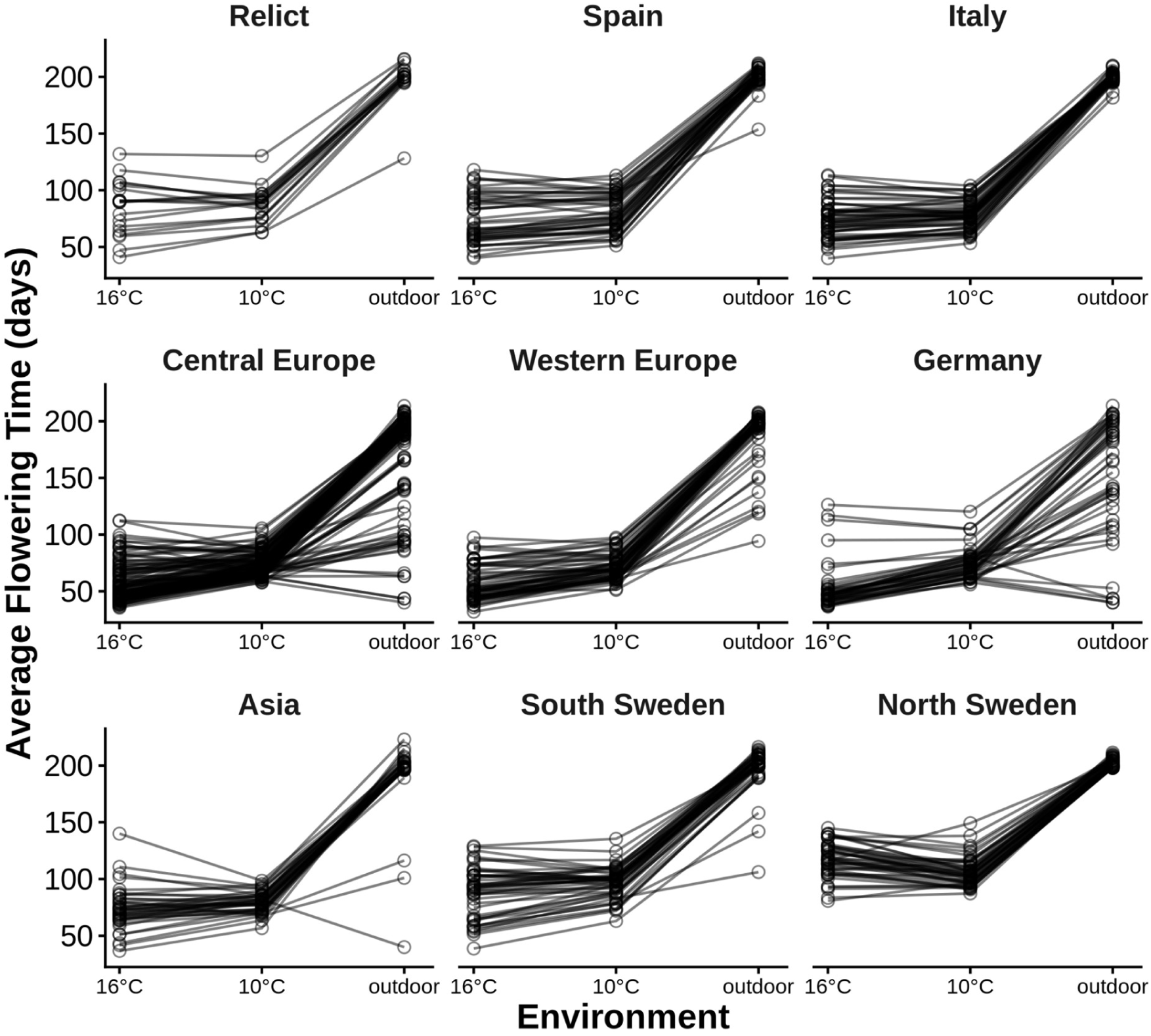
Gene-by-Environment reaction norms comparing flowering time between the 16°C growth chambers, 10°C growth chambers, and the overwinter outdoor common garden. Each data point represents the accession mean flowering time within a given environment. Accessions are grouped into separate reaction norm plots based on admixture groups as assigned by the 1001 Genomes Consortium (2016).

The results are also suggestive of GxE for flowering time (Fig 2, Fig S2, Table S1), which can be driven by changes in the expression of genetic variation or changes in the rank order of genotypes across environments (Bowman 1972). The non-parallel reaction norm slopes between the growth chambers and the outdoor common garden indicate that accessions are responding differently to the different environmental conditions (Fig 2). The Spearman’s rank correlations between the growth chambers and the outdoor common garden was lower (⍴_outdoor, 10°C_ (477) = 0.55, adjusted *p*_outdoor, 10°C_ < 0.001, 95% CI_outdoor, 10°C_ = 0.47, 0.62; ⍴_outdoor, 16°C_(477) = 0.63, adjusted *p*_outdoor, 16°C_ < 0.001, 95% CI_outdoor, 16°C_ = 0.56, 0.69; Fig S2) than the Spearman’s rank correlation between the two growth chambers (⍴_10°C, 16°C_(477) = 0.87, adjusted *p*_10°C, 16°C_ < 0.001, 95% CI_10°C, 16°C_ = 0.85, 0.90; Fig S2). The strong rank correlation between the two growth chambers suggests that the genetic basis of flowering time is mostly conserved between the 10°C and 16°C growth chambers. Accessions that flowered early in the growth chambers tended to flower early in the outdoor common garden (Fig S2). However, some of the last accessions to flower in the growth chambers were also some of the earliest to flower in the outdoor common garden (Fig S2). The moderate rank correlations between the growth chambers and the overwintering common garden suggests there is some conservation for the genetic basis of flowering but there is a considerable amount of GxE for flowering time between the growth conditions.

The extent of crossing reaction norms we observed differed between admixture groups. There was a larger range in reaction norm slopes between the outdoor common garden and the growth chambers for accessions from the Central Europe, Western Europe, and Germany admixture groups. Whereas, accessions from the Spain, Italy-Balkan-Caucausus, and North Sweden admixture groups flowered within a very narrow period of time within the outdoor common garden. Flowering time for all admixture groups has a high rank correlation between the two growth chamber conditions that ranges from 0.65 to 0.95 (Table S1). In contrast, flowering time for most admixture groups has a significant rank correlation between the growth chambers and the outdoor common garden albeit lower compared to just between the two growth chambers (Table S1). The rank correlations for the Asian admixture groups were not significant between the outdoor common garden and the 10°C and 16°C growth chambers. Rank correlations between the outdoor common garden and 16°C for the Western Europe and North Sweden admixture groups were not significant after correcting for multiple comparison testing and the 95% confidence intervals overlapped with zero. The results indicate that flowering time between the growth chambers and outdoor common garden is not genetically correlated across environments for the Western Europe, North Sweden, and Asia admixture groups which is suggestive of GxE due to changes in rank order. However, the lower rank correlations for the Asia and North Sweden admixture groups may be due to a reduction in variance in the overwintering common garden since accessions are more likely to switch ranks when flowering in a narrow period (Table S1, Fig 2).

### Flowering time large-effect loci are condition-specific

To identify how the underlying genetic basis of flowering time is contributing to the observed differences in flowering time between growth conditions, we conducted a multivariate GWAS.

The multivariate GWAS has 20 peaks that surpassed the FDR 0.05 significance threshold, of which only two peaks (peak 1 and 3) surpassed a Bonferroni significance threshold (Fig 3a). None of the genetic variants that were significant in the multivariate GWAS were significant across all three growth conditions (Fig 3). Only three peaks are shared between 10°C and 16°C growth chambers (peaks 1, 3, 20 in Fig 3). Although peak 6 only passes the FDR 0.05 significance threshold in 10°C univariate GWAS, the peak falls just short of the significance threshold in the 16°C growth chamber (Fig 3c-d). 14 peaks were environment specific, 11 of which are specific to the outdoor common garden (peaks 4-5, 7-8, 10-13, 15-16, 19; Fig 3b). Peak 2 is specific to the 10°C growth chamber and peak 9 only present in the 16°C growth chamber. Peaks 14, 17, and 18 are only significant in the multivariate GWAS and not in any of the univariate models.

**Figure 3.**
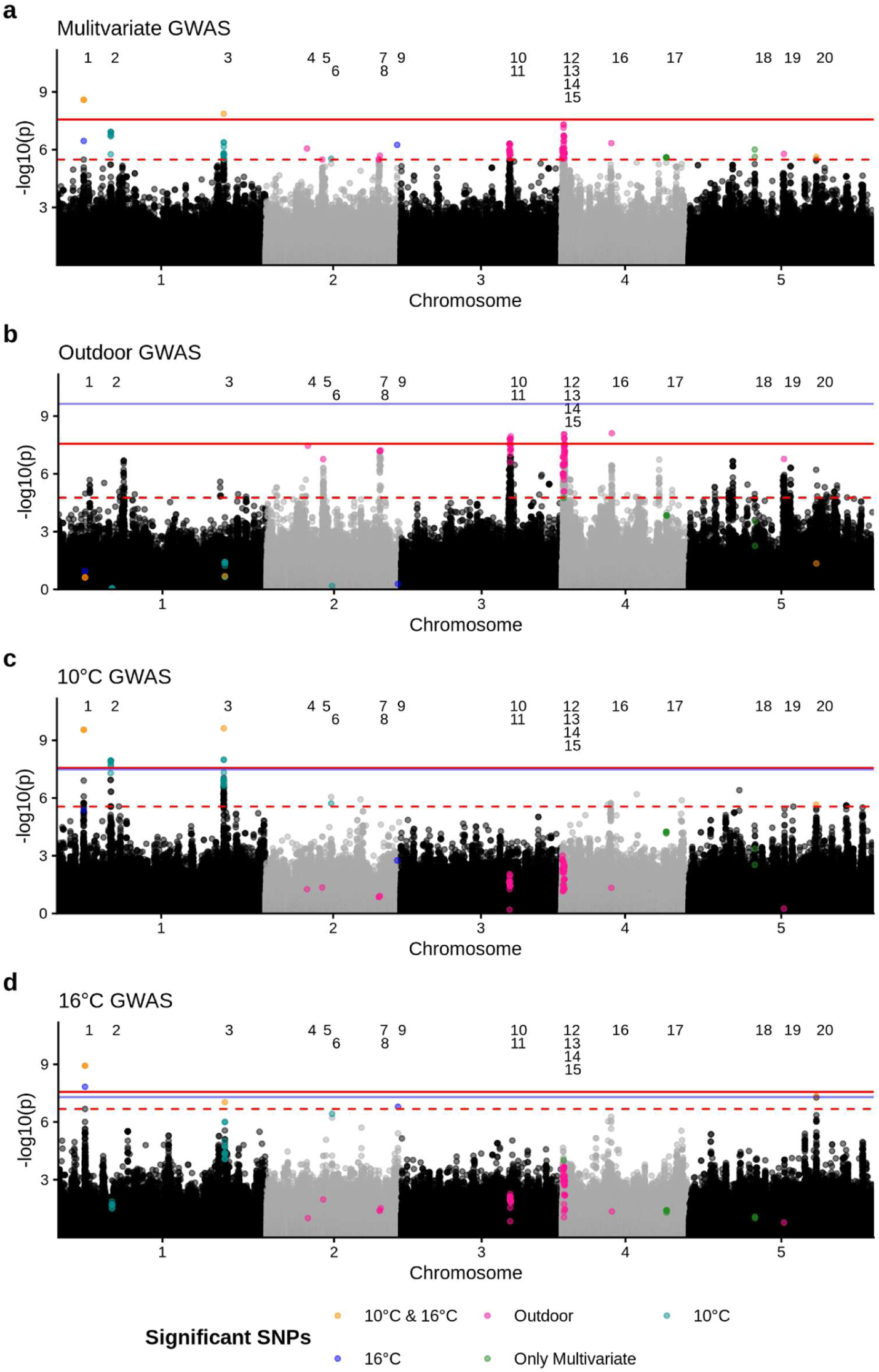
Manhattan plots for flowering time from the **a)** multivariate GWAS, **b)** univariate outdoor common garden GWAS, **c)** univariate 10°C growth chamber GWAS, and **d)** univariate 16°C growth chamber GWAS. The red dashed line represents the largest p-value that surpasses the FDR 0.05 significance threshold. The red solid line is the Bonferroni significance threshold. The blue solid line is the permutation based threshold. Colored data points show SNPs that were significant in the multivariate GWAS and were also significant in at least one of the univariate GWAS. Significant SNPs are grouped into peaks 1-20 labeled at the top of each Manhattan plot.

We identified multiple flowering time candidate genes within 10 Kb of the multivariate GWAS peaks (Table 1). Peaks specific to either one or both of the growth chamber environments included the well known flowering time candidate genes *FLOWERING LOCUS T* (*FT*; AT1G65480), *SHORT VEGETATIVE PHASE* (*SVP*; AT2G22540), and *DAYS OF GERMINATION 1* (*DOG1*; AT5G45830). We also found flowering time candidate genes in multiple peaks specific to the outdoor common garden such as *CENTER CITY* (*CCT*; AT4G00450), *CORONATINE INSENSITIVE 1* (*COI1*; AT2G39940), *NITRILASE 1* (*NIT1*; AT3G44310), *NITRILASE 2* (*NIT2*; AT3G44300), and *TRANSCRIPTION FACTOR JUMONJI DOMAIN-CODING PROTEIN* (*JMJ27*; AT4G00990). Moreover, peaks 7, 12, and 15 included genes *BOTRYTI-INDUCED KINASE 1* (*BIK1*; AT2G39660), *FACTOR OF DNA METHYLATION 2* (*FDM2*; AT4G00380) and *LHP1-INTERACTING FACTOR 2* (*LIF2*; AT4G00830) that have SNPs in exon regions which are significantly associated with flowering time in the multivariate GWAS. Peak 14 that included the well known flowering time gene *FRIGIDA* (*FRI*; AT4G00650) was significant only in the multivariate GWAS but the peak is within a 260 Kb genomic region with multiple peaks (peaks 12, 13, and 15) that are significant in the outdoor common garden. Peak 18 that was significant in the multivariate GWAS but not in any of the condition-specific univariate GWAS, also included candidate gene *GIBBERELLIN INSENSITIVE DWARF 1C* (*GID1C*; AT5G27320). There are eight GWAS peaks with no known flowering time genes. Four GWAS peaks (peaks 5, 10, 16, and 19) were specific to the outdoor common garden. The growth chambers both had one condition-specific peak (peak 2 in 10°C and peak 9 in 16°C) and one shared peak (peak 1) with no candidate flowering time genes. Peak 1 was one of the only peaks that surpassed the Bonferroni threshold in the multivariate GWAS. Additionally, peak 17 that was only significant in the multivariate GWAS and not any of the condition specific univariate GWAS also included no flowering time candidate genes.

**Table 1.**
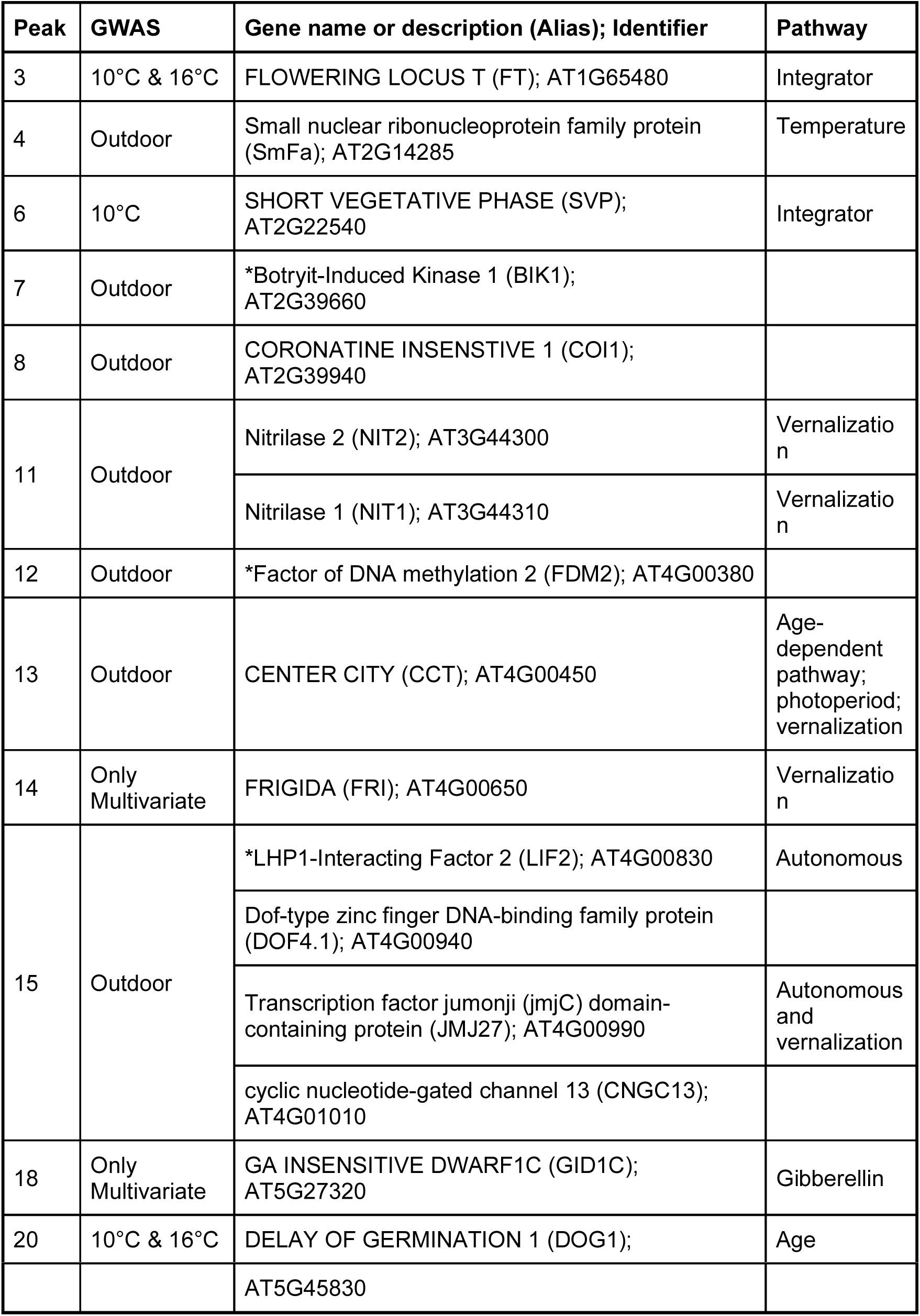
Flowering time candidate genes within 10 kb of GWAS peaks. The full gene list including non-candidate genes can be found in Dataset S4. An * indicates candidate genes with significant SNPs found in genic regions.

| Peak | GWAS | Gene name or description (Alias); Identifier | Pathway |
|---|---|---|---|
| 3 | 10°C & 16°C | FLOWERING LOCUS T (FT); AT1G65480 | Integrator |
| 4 | Outdoor | Small nuclear ribonucleoprotein family protein (SmFa); AT2G14285 | Temperature |
| 6 | 10°C | SHORT VEGETATIVE PHASE (SVP); AT2G22540 | Integrator |
| 7 | Outdoor | *Botryit-Induced Kinase 1 (BIK1); AT2G39660 |  |
| 8 | Outdoor | CORONATINE INSENSTIVE 1 (COI1); AT2G39940 |  |
| 11 | Outdoor | Nitrilase 2 (NIT2); AT3G44300 | Vernalization |
|  |  | Nitrilase 1 (NIT1); AT3G44310 | Vernalization |
| 12 | Outdoor | *Factor of DNA methylation 2 (FDM2); AT4G00380 |  |
| 13 | Outdoor | CENTER CITY (CCT); AT4G00450 | Age-dependent pathway; photoperiod; vernalization |
| 14 | Only Multivariate | FRIGIDA (FRI); AT4G00650 | Vernalization |
| 15 | Outdoor | *LHP1-Interacting Factor 2 (LIF2); AT4G00830 | Autonomous |
|  |  | Dof-type zinc finger DNA-binding family protein (DOF4.1); AT4G00940 |  |
|  |  | Transcription factor jumonji (jmjC) domain-containing protein (JMJ27); AT4G00990 | Autonomous and vernalization |
|  |  | cyclic nucleotide-gated channel 13 (CNGC13); AT4G01010 |  |
| 18 | Only Multivariate | GA INSENSITIVE DWARF1C (GID1C); AT5G27320 | Gibberellin |
| 20 | 10°C & 16°C | DELAY OF GERMINATION 1 (DOG1); | Age |
|  |  | AT5G45830 |  |

Given that loci with smaller effect sizes that are still contributing to differences in flowering time are less likely to surpass stringent GWAS significance thresholds, we next compared the effect sizes of the top 1% of the most significant SNPs (n = 18,339) from the multivariate GWAS (Fig 4, Table S2). Even though using the top 1% of SNPs increases the chances of false positives, the 1% of genes should be enriched for true associations compared to the rest of the genome. Effect sizes were more correlated between the two growth chambers than between the outdoor common garden and the growth chambers (Pearson’s correlation: r_10°C, 16°C_ = 0.80, adjusted *p*_10°C, 16°C_ < 0.001; r_outdoor, 10°C_ = 0.38, adjusted *p*_outdoor, 10°C_ < 0.001; r_outdoor, 16°C_ = 0.49, adjusted *p*_outdoor, 16°C_ < 0.001; degrees of freedom = 18,337). We observed similar correlations when we calculated rank correlations that should buffer against overestimated effect sizes for the most significant SNPs (Spearman’s rank correlation: ⍴_outdoor, 10°C_ = 0.39, adjusted *p*_outdoor, 10°C_ < 0.001; ⍴_outdoor, 16°C_ = 0.50, adjusted *p*_outdoor, 16°C_ < 0.001; ⍴_10°C, 16°C_ = 0.81, adjusted *p*_10°C, 16°C_ < 0.001; degrees of freedom = 18,337). Correlations for the top 1% of SNPs were higher than correlations for 1% randomly sampled SNPs which suggests that the top 1% of SNPs are more similarly contributing to flowering time compared to the genome-wide average (Table S2). However, the rank correlations of SNP effect sizes for the top 1% of SNPs were lower than for rank correlations estimated with phenotypes (Fig S2, Fig 4, Table S2). Together, the results suggest that the most significant SNPs in the multivariate GWAS have condition-specific effects in the outdoor common garden but have similar effects on flowering time in the two growth chamber conditions.

**Figure 4.**
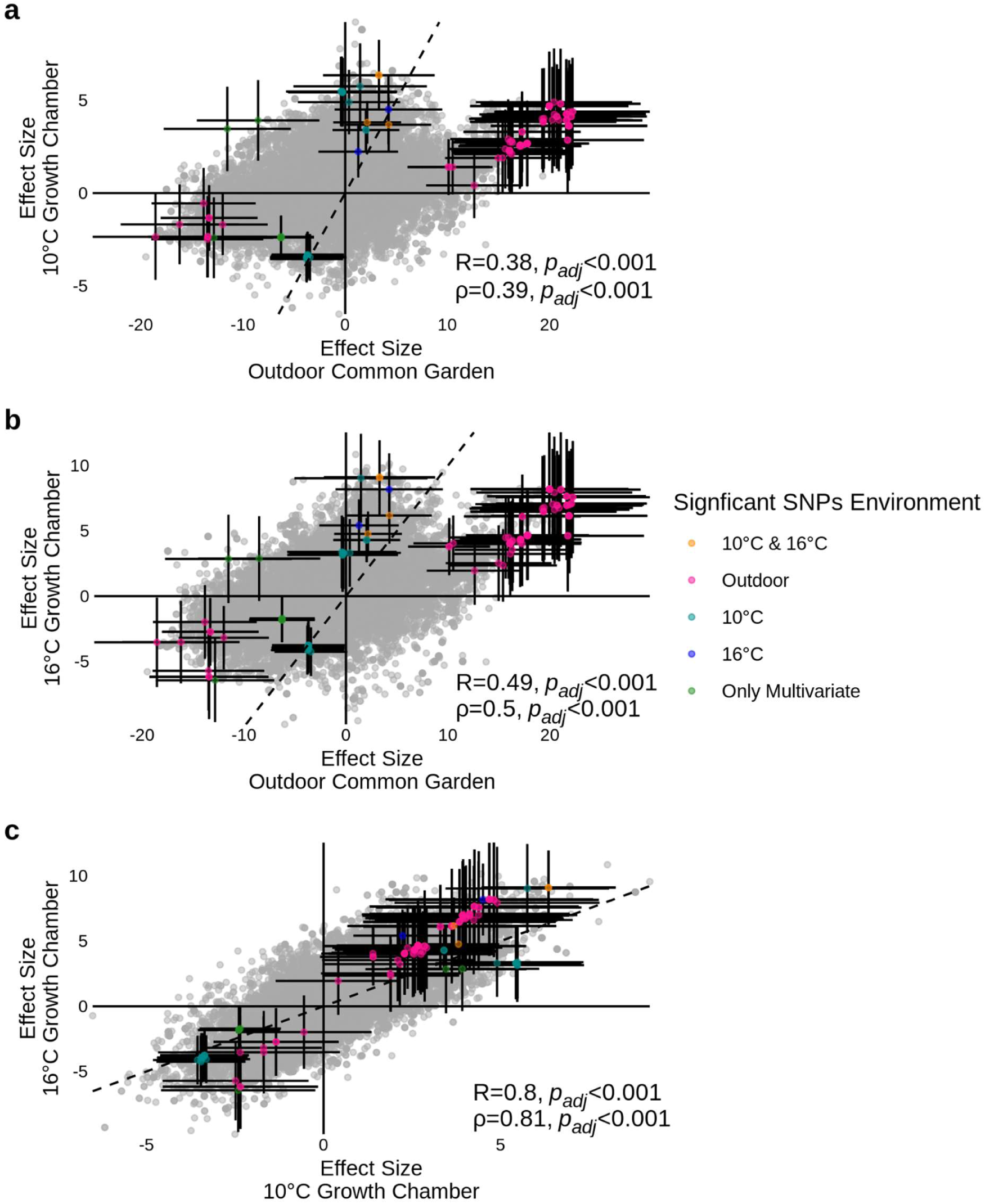
Scatter plots comparing the effect size from the condition-specific univariate GWAS for most significant (top 1%) of SNPs identified in the multivariate GWAS between **a)** the 10°C growth chamber and the outdoor common garden, **b)** the 16°C growth chamber and the outdoor common garden, and **c)** the 10°C and 16°C growth chambers.The dashed black line represents the 1:1 line. Colored data points show SNPs that were significant in the multivariate GWAS and were also significant in at least one of the univariate GWAS as shown in Fig 3. Error bars for colored data points represent 95% confidence intervals (±1.96*SE). The Pearson’s correlation and Spearman’s rank correlation statistics and adjusted p-values are reported in the bottom corner for each plot.

We next determined if large effect loci were contributing to GxE through antagonistic pleiotropy, allelic differential sensitivity, and conditional neutrality. SNPs with non-overlapping 95% confidence intervals between the outdoor common garden and the growth chambers provide further support for SNPxE underlying the observed GxE for flowering time (Fig 4a-b).

The significant SNPs from the multivariate GWAS that were also significant in the outdoor common garden had different effects (no effect, effects in opposite directions, or same direction but different magnitude) in at least one if not both of the growth chambers (Table S3). However, we found no evidence of SNPxE between the two growth chambers as the effect size 95% confidence intervals overlapped (Fig 4c, Fig S3, Table S3). All of the significant SNPs from the multivariate GWAS that were significant in the growth chambers had a similar effect size in the outdoor common garden (Fig 4a-b, Table S3). Therefore, the loci explaining the most amount of variation in the growth chambers have a similar effect in the outdoor common garden, but loci with the largest impact on flowering time in the outdoor common garden appear condition-specific. The most common SNPxE we observed between the outdoor common garden and the growth chambers was SNPs with allelic differential sensitivity that have effects in the same direction but of a smaller magnitude in the growth chambers compared to the outdoor common garden (Fig S3). We found fewer instances of SNPs that were conditionally neutral that had an effect in the outdoor common garden but not in at least one the growth chambers (Fig S3). We only found one GWAS peak (peak 17) with evidence for antagonistic pleiotropy between the 10°C growth chamber and the outdoor common garden but the locus was neutral in the 16°C growth chamber (Table S3).

### Polygenicity of flowering time differs between growth conditions

We next explored whether there was GxE for the polygenic architecture of flowering time. To do so, we used BSLMM GWAS from GEMMA to estimate the proportion of phenotypic variation explained by genotype (PVE), the portion of genetic variance explained by large effect loci (PGE), and the number of large effect loci affecting the trait (n_ɣ_). Overall flowering time was highly heritable across all three conditions but PVE was lower in the outdoor common garden (mean PVE_outdoor_= 0.71) compared to both of the growth chambers (mean PVE_10°C_= 0.87 and mean PVE_16°C_= 0.94; Fig 5a). Polygenic traits with an infinitesimal architecture should have a PGE close to 0, oligogenic traits should approach 1, and traits with an omnigenic architecture should be in between. Flowering time in the outdoor common garden follows are more oligogenic genetic architecture (mean PGE_outdoor_= 0.93), whereas flowering time in the growth chambers had more of an omnigenic architecture (mean PGE_10°C_= 0.54 and mean PGE_16°C_= 0.52; Fig 5b). Although the amount of genetic variance explained by large effect loci differs between the growth chambers and the outdoor common garden, the number of large effect loci are similar across all three growth conditions (mean n_ɣ outdoor_ = 30.06, mean n_ɣ 10°C_ = 20.05, mean n_ɣ 16°C_ = 45.69; Fig 5c).

**Figure 5.**
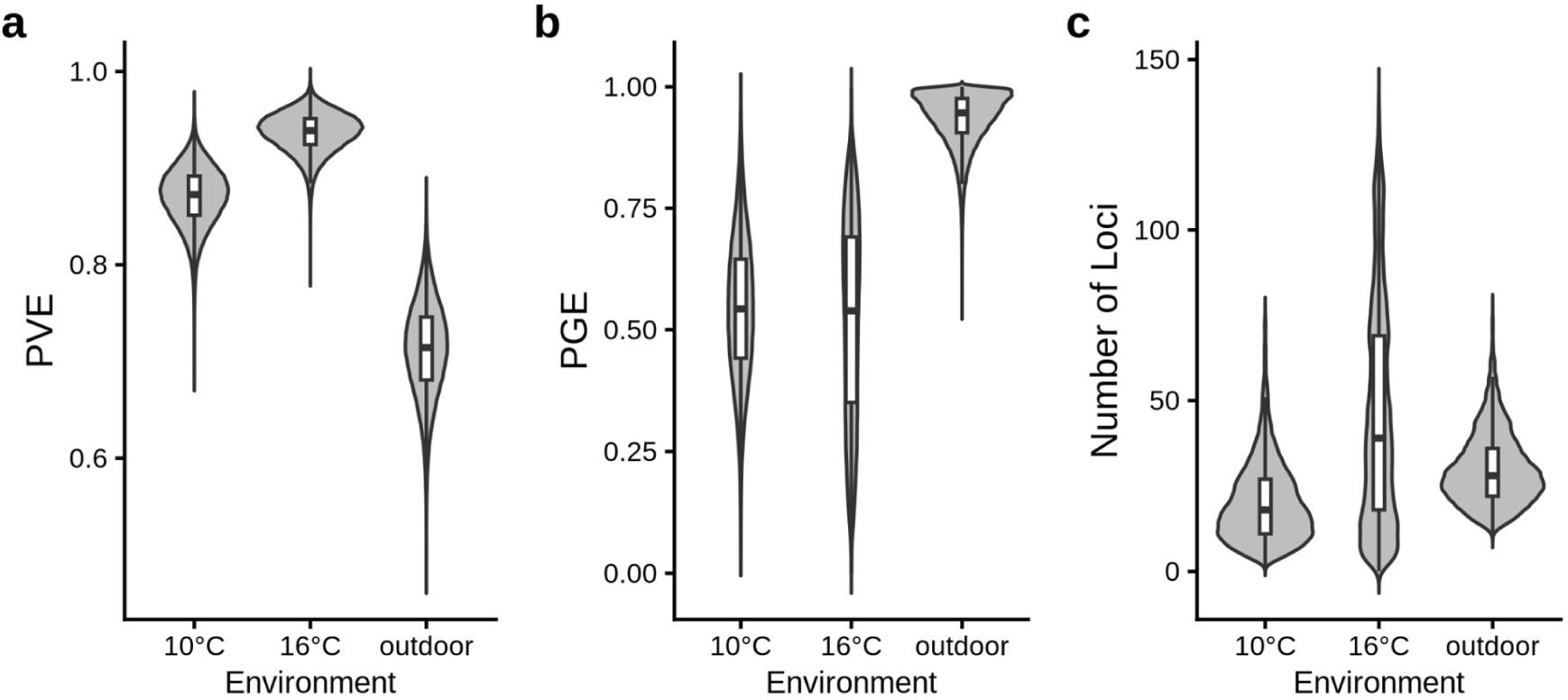
BSLMM posterior distributions for the **a)** proportion of phenotype variance explained by genotype (PVE), **b)** proportion of genetic variance explained by large effect loci (PGE), and **c)** estimated number of large effect loci for flowering time in the 10°C growth chamber, 16°C growth chamber, and outdoor common garden.

### Assessing the robustness of GWAS inferences

We next explored whether the skewed distribution of flowering time in the outdoor common garden affected our inferences of GxE and SNPxE interactions. On the one hand, LMM and BSLMM GWAS models assume normally distributed residuals, and skewed distributions and outliers can affect inferences (e.g., due to rarity disequilibrium). On the other hand, the outdoor common garden experiment clearly showed differences in the mean, variance, and distribution of flowering time; these differences were the *a priori* object of study, and transformations to eliminate them might eliminate the signal of interest. We first calculated a proxy for genome-wide residuals for the outdoor common garden flowering time data to determine if we were violating the assumption of normally distributed residuals. The genome-wide residuals were not normally distributed (Shapiro-Wilk Normality Test: W = 0.856, *p* < 0.001; Fig S7c-d). We tried to assess the effects of non-normal residuals on our GxE and SNPxE inferences by 1) removing individuals with residuals that severely deviated from normality, 2) calculating a permutation based threshold, and 3) transforming the outdoor overwinter phenotype data.

We rerun the univariate LMM and BSLMM GWAS after excluding accessions that most severely violated the assumption of residual normality (Fig S7d). Many of the significant SNPs identified in the multivariate GWAS with the full dataset were either significant or in close proximity to GWAS peaks in the univariate GWAS with the filtered dataset (Fig S9). Overall the BSLMM results were comparable between the full dataset and the filtered dataset for flowering time in the two growth chambers (mean PVE_10°C filtered_= 0.86, and mean PVE_16°C filtered_= 0.92; mean PGE_10°C filtered_= 0.59, and mean PGE_16°C filtered_= 0.46; Fig S10). The proportion of phenotypic variance explained by genotype for the outdoor common garden increased after filtering (mean PVE_outdoor filtered_= 0.93; Fig S10a). The proportion of genetic variance explained by large effect loci decreased for the outdoor common garden but the mean PGE for the outdoor common garden was still higher than growth chambers for the filtered dataset (mean PGE_outdoor filtered_= 0.77; Fig S10b). Although the estimated number of large effect loci decrease for all growth conditions, the overall trend was similar to the full dataset (mean n_ɣ outdoor filtered_ = 21.28, mean n_ɣ 10°C filtered_ = 17.63, mean n_ɣ 16°C filtered_ = 30.14; Fig S10c). Although the PGE decreased in the outdoor common garden, flowering time in the outdoor common garden was still in line with an oligogenic genetic architecture even after filtering. Therefore, our inferences about GxE, SNPxE, and the polygenetic architecture of flowering time after we removed accessions that most severely violated residual normality are mostly consistent with our inferences based on the full dataset.

## Discussion

We used a combination of GWAS approaches to show how the genetic basis of flowering time in *Arabidopsis* changes when grown in an ecologically realistic environment. We found that SNPxE for large effect loci and shifts in the overall polygenic architecture contribute to the extensive GxE we observed for flowering time. Below, we discuss these results in context of *Arabidopsis’* evolutionary history and the known flowering time genetic network. We also consider multiple potential genetic mechanisms that could be underlying GxE for flowering time.

### GxE is consistent with *Arabidopsis’* evolutionary history

The amount of GxE we observed for flowering time across the global set of accessions is in line with other studies that have shown differences in flowering time between field and lab conditions (Brachi et al. 2010; Méndez-Vigo et al. 2013; Stock et al. 2015). We also found that GxE differs between admixture groups. For some admixture groups the variation in flowering time increases in the outdoor common garden such as Central Europe, Western Europe, and Germany (Fig 2). Multiple loss-of-function alleles for genes involved in the vernalization pathway are maintained in central and western European regions, such as *FRI* and *FLOWERING LOCUS C* (*FLC*) that allow individuals to flower without exposure to a prolonged cold period (Johanson et al. 2000; Le Corre et al. 2002; Gazzani et al. 2003). Accessions from the Central European region are thought to be weedy rapid cyclers that can flower in both spring to early summer as well as late summer to fall (Exposito-Alonso 2020). In contrast, other admixture groups such as North Sweden and Italy-Balkan-Causasus have the opposite effect of reduced flowering time variation in the outdoor common garden where all accessions flowered within a very narrow window (Fig 2). Accessions from North Sweden are from the northern part of the species range where colder and longer winters are a more prominent selective force (Ågren and Schemske 2012; Yim et al. 2024). Accessions from the northern part of the species range are thought to have strict vernalization requirements to ensure that individuals germinate in the fall, overwinter as rosettes, and flowering in the spring (Exposito-Alonso 2020). The results are consistent with North Sweden accessions having adapted to prolonged cold.

### Outdoor common garden reveals genes across multiple regulatory pathways

We found no overlapping significant large-effect loci between the outdoor common garden and the growth chambers (Fig 3). The large-effect loci associated with flowering time in the growth chambers are in close proximity to genes *FT*, *SVP*, and *DOG1* (Table 1). All three of these genes were previously identified as being associated with flowering time in the larger set of 1003 accessions in the 10°C growth chamber (1001 Genomes Consortium 2016). *FT* acts downstream of multiple regulatory flowering time pathways and is often referred to as an integrator gene in the flowering time genetic network (Abe et al. 2005; Wigge et al. 2005; Schwartz et al. 2009). *SVP* together with *FLC* negatively regulates *FT* (Li et al. 2008). The gene *DOG1* regulates the levels of microRNAs that affect the time of the transition between vegetative and reproductive stages in *Arabidopsis* (Huo et al. 2016). Interestingly, we did not detect a GWAS peak in close proximity to *DOG1* in the overwintering common garden even though some of the earliest work implicating *DOG1* as a flowering time gene came from experiments done in the field (Chiang et al. 2013).

We were also able to identify multiple well-known flowering time candidate genes in the overwintering common garden including *CCT, LIF2*, *JMJ27*, *CNGC13*, *NIT1*, and *NIT2* (Table 1). The genes *CCT, FRI, JMJ27*, *CNGC13*, and *LIF2* are all located within 250kb of each other on Chromosome 4. Although the GWAS peak with *FRI* is only significant in the multivariate GWAS, the peak is nested within a region that has multiple peaks that are significant in the outdoor common garden. *FRI* is a well known flowering time gene in the vernalization pathway that promotes *FLC* to repress flowering until after a prolonged cold period (Michaels and Amasino 1999). There are multiple *FRI* alleles found within the 479 accessions (Zhang and Jiménez-Gómez 2020) and variation in *FRI* has previously been shown to play an important role in naturally occurring variation in flowering time (Johanson et al. 2000; Caicedo et al. 2004; Stinchcombe et al. 2004; Atwell et al. 2010; Fournier-Level et al. 2022). Furthermore, *FRI* has been shown to be involved in GxE between field and glasshouse growth conditions (Méndez-Vigo et al. 2013). *CCT* regulates multiple flowering time genes including *FT*, *FLC*, and *SOC1* (Imura et al. 2012). Similarly, *LIF2* promotes *FLC* in short and long days but *lif2* has no effect on flowering time when grown in continuous light (Latrasse et al. 2011). Although the effect of *LIF2* on flowering has been shown using mutant screens, the gene has not previously been implicated in affecting naturally occurring flowering time variation. *JMJ27* is also thought to regulate multiple flowering time genes including *FLC* and *CONSTANS* (*CO*) and is involved in the vernalization pathway (Dutta et al. 2017). Interestingly, a recent study testing a conditional GWAS approach using the 10°C growth chamber identified a new hit that overlaps with peak 15 and experimentally validated that both *JMJ27* and *CNGC13* resulted in earlier flowering time (Bellis et al. 2026). Given that *CCT, FRI, JMJ27*, *CNGC13*, and *LIF2* are all within the same genomic region, it is challenging to determine which candidate gene or genes are driving the peak (Atwell et al. 2010; 1001 Genomes Consortium 2016). Additionally, GWAS peak 9 on Chromosome 3 includes a tandem array of *NIT2-NIT1-NIT3*. Together *NIT1-3* have been shown to repress *MADS AFFECTING FLOWERING 4* (*MAF4*) which is a paralog of *FLC* (Yang et al. 2022). Although *NIT1-3* triple mutants show an effect on flowering time in both short and long days, flowering time is unaffected in constant light conditions which indicates the effects of these genes can be condition specific (Yang et al. 2022). Even though genetic manipulation studies show a clear effect on flowering time, *NIT2-NIT1-NIT3* has not previously been identified as affecting flowering time in naturally occurring variation. Therefore, our results provide additional ecological insight for flowering time genes such as *NIT1*, *NIT2*, and *LIF2*.

We identified three multivariate GWAS peaks (peaks 14, 17, and 18) that were not significant in any of the condition-specific univariate GWAS (Fig 3). One of the strengths of phenotyping individuals across multiple environments is that multivariate GWAS can have increased power to detect loci affecting a phenotype across multiple environments compared to univariate approaches (Galesloot et al. 2014). *GIBBERELLIN INSENSITIVE DWARF1C* (*GID1C*), a gene involved in the gibberellic acid flowering time pathway, is found in close proximity to GWAS peak 18 (Fig 3). *GID1C* has not previously been identified as a candidate gene in a GWAS, likely due to its potential functional redundancy with *GID1B* and *GID1A* (Griffiths et al. 2006). However, a transcription-wide association study of 690 *Arabidopsis* accessions study showed that variation in transcription levels of *GID1C* was associated with flowering time (Li et al. 2021). The multivariate GWAS results suggest *GID1C* could be affecting flowering time, although its effects might be too subtle or weak to detect in a univariate GWAS, especially if there is some functional redundancy with *GID1A* and *GID1B*.

We identified multiple GWAS peaks that were significantly associated with flowering time but did not include known flowering time genes. There were a total of eight GWAS hits with no known flowering time genes, four of which were specific to the outdoor common garden, three were specific to either one or both growth chambers, and one hit was only significant in the outdoor common garden experiment. One interpretation is that these hits include novel flowering time genes that have not yet been identified. However, an alternative explanation is that the hits with no known flowering time genes could be false positives. Therefore, these eight genomic regions require further experimental validation ideally involving genetic manipulation and phenotyping in a variety of growth conditions - especially for regions that were specific to the outdoor common garden or only identified in the multivariate GWAS.

### Genetic mechanisms underlying GxE

There are multiple possible genetic mechanisms that can give rise to GxE, such as conditional neutrality, allelic differential sensitivity, and antagonistic pleiotropy. Conditional neutrality or allelic differential sensitivity is the most straightforward explanation for condition specific GWAS hits, especially for genes that respond to external stimuli that are present in the overwintering common garden but absent in the growth chambers. Therefore, it is not surprising that most of the SNPxE we observed between the outdoor common garden were from SNPs in close proximity to candidate genes in the temperature, vernalization, and photoperiod flowering time pathways with either a smaller effect or no effect on flowering time in the growth chambers. However, comparing effect sizes between GWAS should be done with caution since effect sizes of the most significant SNPs are likely to be overestimated - a statistical limitation of GWAS often referred to as the winner’s curse (Göring et al. 2001). Additionally, GWAS methods have a bias against detecting loci with smaller effect sizes. It is possible that some SNPs that we consider to be conditionally neutral in reality have an effect on flowering time in both environments but due to the statistical limitations of current GWAS models or sample size we were unable to detect an effect. Both the moderately correlated effect sizes for the top 1% of multivariate GWAS SNPs and some of the SNPs with overlapping 95% confidence intervals for effect sizes between environments suggests there is a partially shared genetic basis of flowering time between the outdoor common garden (Fig 4, Fig S3, Table S2-3). We only found one locus, GWAS peak 18 which includes candidate gene *GID1C*, to have antagonistic pleiotropic effects on flowering time between the 10°C growth chamber and outdoor common garden but the locus is neutral in the 16°C growth chamber. *GID1C* could thus be important for naturally occurring variation in flowering time with different effects across multiple environments.

## Conclusions

Our work highlights the importance and challenges of measuring complex traits in conditions that better mimic those found in nature. We show how the effects of individual loci, alternative regulatory pathways, and the overall polygenic architecture contribute to differences in flowering time phenotypic distributions between an ecologically realistic environment and standard lab conditions. Our findings show that inferences about genetic architecture, evolution, and the maintenance of adaptive phenotypic variation may be critically dependent on how closely the experimental environment mimics natural variation.

## Supporting information

Supplement including: supplementary methods, supplementary Note1, Supplementary Figure S1-10, Supplementary Tables S1-4

## Author Contributions

ALP: Conceptualization, methodology, formal analysis, investigation, data curation, writing - original draft, visualization, funding acquisition

JRS: Conceptualization, resources, writing - review & editing, supervision, funding acquisition

## Acknowledgments

We thank Aneil Agrawal, Stephen Wright, Megan Bontrager, and members of the Stinchcombe Lab for advice on the project. Undergraduates Katherine Leung, Jessica Underwood, Yasmin Rabiee, Aidan Godin, Grace Walker Mitchell, Ishani Sharma, Alice Xiao, Kieran Guimond, Katie Monat, Anya Gopaul, Claire Abbasi, Michelle Wong, Trisha Leigh Malate, Emma Zongaro, and Ellie Hector assisted with transplanting and data collection. We thank Tom Gludovacz and Alice DesRoches for horticultural support. Our research was enabled by the computational resources and support provided from Compute Ontario and the Digital Research Alliance of Canada.

