## Supplement including: supplementary methods, supplementary Note1, Supplementary Figure S1-10, Supplementary Tables S1-4 for "GxE and changes to the genetic architecture of flowering time in an overwintering environment"

**This PDF file includes:**

Supplementary Methods
Supplementary Note 1
Supplementary Figures S1-10
Supplementary Tables S1-S4

### Supplementary Methods

Germination conditions and transplanting

We stratified seeds for 7 days at 4°C in 0.15% agar. After stratification, on September 29th 2022, we sowed individuals seeds onto Promix soil in 3-inch coco coir with eight replicates per accession. The greenhouse was set to 20-23°C daytime temperature, 15-20°C night temperature, and an 11 hour photoperiod. Germination domes were used for the first 17 days to ensure high humidity was maintained during germination. To avoid overheating on sunny days, germination domes were propped open from 11am to 4pm. Individuals were misted daily except when germination domes were propped open then they were misted twice a day. Individuals were acclimatized to colder conditions on October 19th 2022 before transplanting by setting the greenhouse to 12-16°C daytime and 6-9°C night temperature with no supplemental light. Each block in the outdoor common garden included one replicate per accession. We transplanted ungerminated replicates only if there were less than six individuals that had germinated out of the eight originally sown. Transplanting took place over 2 days (October 24-25, 2022); blocks 1-3 were transplanted on the first day and blocks 4-6 were transplanted on the second day.

Temperature tracking for rooftop common garden

To measure the natural temperature variation experienced by the plants (Dataset S3), on December 6th 2022, we deployed DS1921G-F5# Thermochron iButton data loggers within DS9107 iButton Capsules that recorded the temperature every four hours. Within each experimental bed, iButtons were placed in two positions, in between the middle row of plants at both the north and south sides of the beds. We placed iButtons at both the surface and approximately 3 cm below the surface. iButtons on the surface were removed on April 15th 2023 to stop squirrels from coming into the beds to move the iButtons. For temperature data before the iButtons were deployed, from October 24th 2022 - December 6th 2022, we used temperature data collected by the University of Toronto growth facilities from the same building as the rooftop common garden. Temperature data was processed using R package lubridate v1.9.4 (Grolemund and Wickham 2011).

Cleaning phenotype data

Data for all phenotypes were formatted and merged using a custom R script. We then checked for any data inconsistencies. For example, we checked if any individuals had been recorded to have flowered before they bolted. We either programmatically fixed any errors that were verifiable or excluded data that we could not confidently verify. We removed accessions that only had flowering time data for a single replicate. After cleaning the dataset and removing all individuals that did not survive until flowering, we had 479 accessions in the finalized dataset (Dataset S1). For the downstream analyses, we used the accession mean across all blocks.

Visualisations and data analyses in R:

Visualisations were performed in R 4.4.0 (R Core Team 2024) using Rstudio v2026.10.999.999 (Posit Team 2024). Figures were generated using ggplot2 v4.0.0 (Wickham 2016), ggvenn v0.1.19 (Yan 2025), ggrepel v0.9.6 (Slowikowski 2024), ggpubr v1.0.0 (Kassambara 2023), scales v1.4.0 (Wickham et al. 2025), ggforce v0.5.0 (Pedersen 2025a), and patchwork v1.3.2 (Pedersen 2025b). We used packages readxl v1.4.5 (Wickham and Bryan 2025), dplyr v1.2.1 (Wickham et al. 2023), GenomicRanges v1.58.0 (Lawrence et al. 2013), tidyr v1.3.2 (Wickham et al. 2024), data.table v1.16.0 (Barrett et al.

2024), rlang v1.3.0 (Henry and Wickham 2026), and conflicted v1.2.0 (Wickham 2023) to import and prepare data for visualisations and analyses. R packages used to run statistical analyses included lme4 v1.1-37 (Bates et al. 2015), lmerTest v3.2-1 (Kuznetsova et al. 2017), boot v1.3-32 (Canty and Ripley 2025), and car v3.1-3 (Fox and Weisberg 2019).

**Supplementary Note 1: Evaluating the robustness of GWAS inferences with skewed phenotypic distributions**

GWAS, especially for complex traits, are notorious for increased risk of type I and type II error. We note that the left-skewed distribution of flowering time in the overwintering common garden further increases the risk of type I error (Fig S7a). Especially since the genome-wide residual proxies calculated with BSLMM are not normally distributed (Fig S7c-d). Skewed phenotypic distributions and outliers can result in spurious associations due the increased occurrence of covariation between lower minor allele frequencies and extreme phenotypes by random chance, sometimes referred to as rarity disequilibrium (Houle and Márquez 2015; Brown and Kelly 2022). However, we believe that the left-skewed distribution has a biological basis and *Arabidopsis* accessions flowering prior to winter by either bypassing vernalization or differing in vernalization requirements are an important aspect of naturally occurring flowering time variation (Bloomer and Dean 2017). We took three approaches to assess the effects of non-normal residuals on our inferences by 1) removing accessions with residuals that severely deviated from normality, 2) calculating a permutation based threshold, and 3) transforming the data. Below we describe how we performed these analyses, the results, as well as the benefits and costs of each approach.

<u>Approach 1: Removing accessions with residuals that severely deviated from normality</u>

Since GWAS models assume that residuals are normally distributed, we removed accessions with genome-wide BSLMM residuals that most strongly deviated from normality. The benefit of this approach is that we are directly removing accessions that violate the assumption of normality but we retained the original phenotypic values of the remaining individuals and the filtered phenotypic distribution still resembles the full dataset. However, we do reduce the sample size which reduces the power and also changes the genetic composition included in the GWAS.

*Methods:*

We removed accessions on either tail of the normalized quantile distribution that diverged from the QQ referline line (Fig S7d). We retained 417 individuals with -1.25 and 2 of the normalized quantile distribution as shown within the two dashed red lines in Fig S7d. We then reran the three LMM univariate GWAS and BSLMM with the filtered subset of accessions for all three growth conditions as described in the main methods (Multivariate GWAS and BSLMM sections).

*Results:*

Many of the significant SNPs identified in the multivariate GWAS with the full dataset were either significant or in close proximity to GWAS peaks in the univariate GWAS with the filtered dataset (Fig S9). Overall the BSLMM results were comparable between the full dataset and the filtered dataset for flowering time in the two growth chambers (mean $PVE_{10°C\ filtered}$= 0.86, and mean $PVE_{16°C\ filtered}$= 0.92; mean $PGE_{10°C\ filtered}$= 0.59, and mean $PGE_{16°C\ filtered}$= 0.46; Fig S10). The proportion of phenotypic variance explained by genotype for the outdoor common garden increased after filtering (mean $PVE_{outdoor\ filtered}$= 0.93; Fig S10a) which makes sense as trait heritability should increase after excluding outlier residuals. The proportion of genetic variance explained by large effect loci decreased for the outdoor common garden but the mean PGE for the outdoor common garden was still higher than growth chambers for the filtered dataset (mean $PGE_{outdoor\ filtered}$= 0.77; Fig S10b). Although the estimated number of large effect loci decrease for all growth conditions, the overall trend was similar to the full dataset (mean $n_{\gamma\ outdoor\ filtered}$ = 21.28, mean $n_{\gamma\ 10°C\ filtered}$ = 17.63, mean $n_{\gamma\ 16°C\ filtered}$ = 30.14; Fig S10c). Although the PGE decreased in the outdoor common garden, flowering time in the outdoor common garden was still in line with an oligogenic genetic architecture even after filtering. Therefore, our inferences about GxE, SNPxE, and the polygenetic architecture of flowering time after we removed accessions that most severely violated residual normality are mostly consistent with our inferences based on the full dataset.

<u>Approach 2: Calculating a permutation based threshold</u>

GWAS typically use Bonferroni or FDR 0.05 significance thresholds, however, neither of which take into account how the phenotypic or genetic distributions may be affecting the risk of Type I error. The benefit of using a permutation based threshold should account for the underlying skewed phenotypic distribution. However, as described below the permutation based threshold we calculated for the outdoor common garden is orders of magnitude more stringent than a Bonferroni correction which alleviates risk of type I error but increases the risk of type II error.

*Methods:*

We conducted 10,000 permutations where we shuffled the phenotype values randomly using the “shuf” bash command. We then conducted a univariate GWAS for the outdoor common garden using the shuffled phenotype values. We saved the most significant p-value from the 10,000 runs from which we used the 5th percentile as the permutation based threshold. Note that this procedure is the GWAS analog of what was developed for QTL mapping by (Churchill and Doerge 1994).

*Results:*

The permutation based threshold for the outdoor common garden was $2.32x10^{-10}$ which is multiple orders of magnitude more stringent than the $2.73x10^{-8}$ Bonferroni threshold and the $1.73x10^{-5}$ FDR 0.05 threshold (Fig 3b). As a result, no SNPs surpassed the permutation based threshold in the outdoor common garden univariate GWAS. The permutation based threshold for the 10°C growth chamber was $3.31x10^{-8}$ and the 16°C growth chamber was $5.06x10^{-8}$, which were both very similar to the Bonferroni threshold.

<u>Approach 3: Transforming the data</u>

We also tried to transform the flowering time phenotypic data in an attempt to normalize the phenotypic distribution. We tried multiple transformations as described below but most transformations did not normalize the outdoor common garden phenotypic distribution except for the rank-based inverse normal transformation. But transforming the overwintering flowering time data using a rank-based inverse normal transformations can reduce power (Beasley et al. 2009). Such dramatic transformations can increase the risk of type II error especially if the skewed distribution has a biological basis. Therefore, transforming the left skewed distribution to a unimodal distribution to fit the statistical model also potentially discards important biological information that were of interest to this study.

*Methods:*

We tried to transform the outdoor common garden flowering time data. We tried multiple transformations including log, reflected log, square root, boxcox, and rank-based inverse normal transformation (Table S4). The boxcox transformation was performed using the car R package. The rank-based inverse normal transformation was performed using the R package RNOmni v1.0.1.2 (McCaw et al. 2020), where ties in rank order are assigned average rank values. The rank-based inverse normal transformation was the only transformation that normalized the outdoor common garden flowering time data (Table S4). We used the rank-based inverse normal transformation for flowering time in all three growth environments and reran the multivariate, condition-specific LMM univariate, and BSLMM GWAS analyses.

*Results:*

The rank-based inverse normal transformations result in reduced power to detect large effect loci across all GWAS analyses (Fig S8; Fig S10). Only one locus surpasses the significance thresholds in the transformed multivariate GWAS (Fig S8a) which corresponds to GWAS peak 3 in the Fig 3. The condition specific univariate GWAS show a reduction in power across all three environments even though the growth chamber flowering time distributions were more normally distributed than the outdoor common garden (Fig S8b-d). There are no significant associations in the transformed outdoor common garden univariate GWAS. Even in the growth chambers, we are no longer able to identify candidate genes such as *SVP* and *DOG1*. The only peak retained in the transformed multivariate GWAS is in close proximity to the well known *FT* flowering time gene. Therefore, *FT* still showed a condition-specific effect on flowering time, suggesting that the genetic basis of the trait differed between growth conditions, regardless of the analysis approach. These results also still indicate the presence of GxE and SNPxE interactions, but we are only able to identify one large effect loci in the growth chambers and none in the outdoor common garden. With the transformed data, the BSLMM results estimate a much lower proportion of genetic variance explained by large effect loci across all three growth conditions (mean $PGE_{outdoor\ transformed}$= 0.26; mean $PGE_{10°C\ transformed}$= 0.24; mean $PGE_{16°C\ transformed}$= 0.31; Fig S10). Since the transformation resulted in reduced power to detect LMM multivariate and univariate GWAS hits, it is not surprising that the polygenic architecture for the transformed data resembles an infinitesimal model (Fig S10). Although using the untransformed data increases the risk of type I error, the transformation appears to also increase type II error across all environments for identifying large effect loci which impacts the ability to accurately infer the polygenetic architecture. Although rank-based inverse normal transformations have commonly been used to normalize phenotype data in GWAS, the transformation can reduce power and may not control type I error (Beasley et al. 2009).

**Figures**

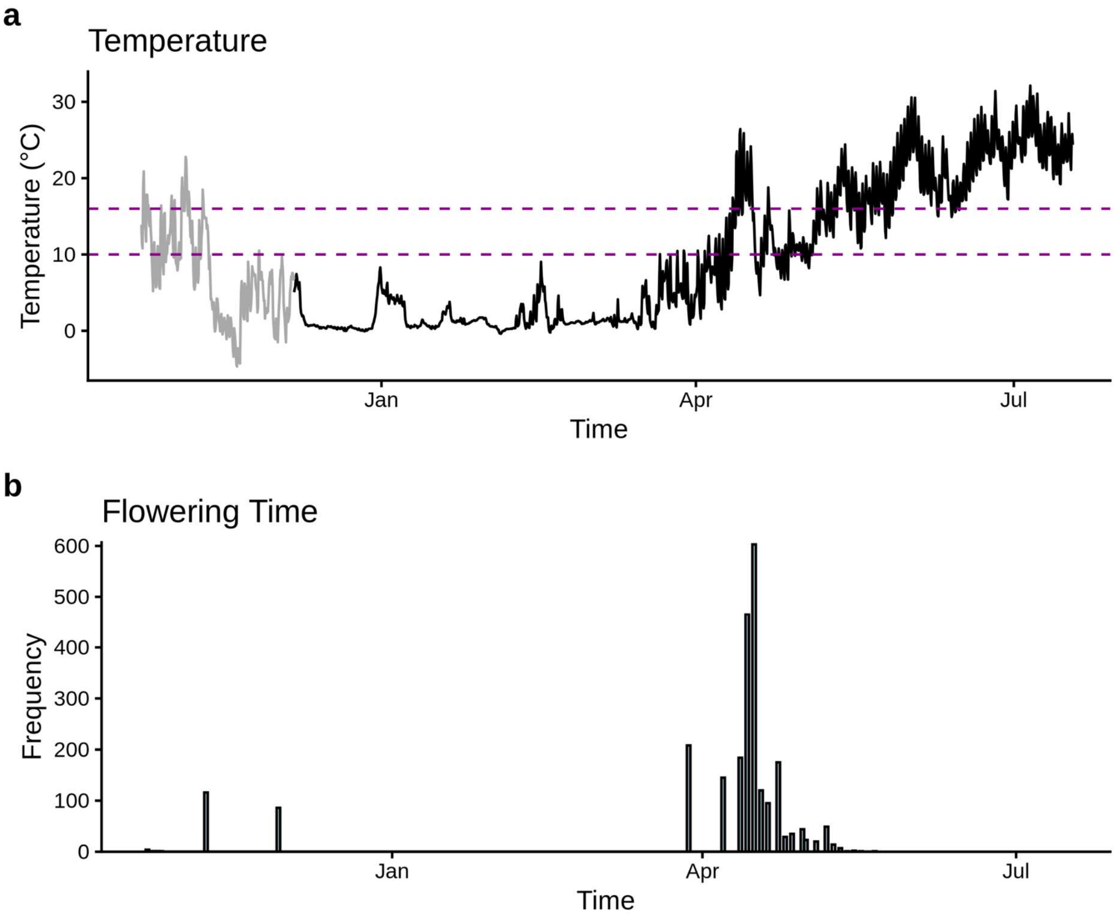


**Figure S1. a)** Temperature in the outdoor common garden. Black line is the average iButton temperatures across all blocks. Grey line is the air temperature before iButtons were deployed into the raised beds. Purple dashed lines show the constant growth chamber temperatures (10°C and 16°C) for the flowering time data from the 1001 Genomes Consortium. **b)** Flowering time for all individuals (n=2443) after transplanting into the outdoor common garden experiment.

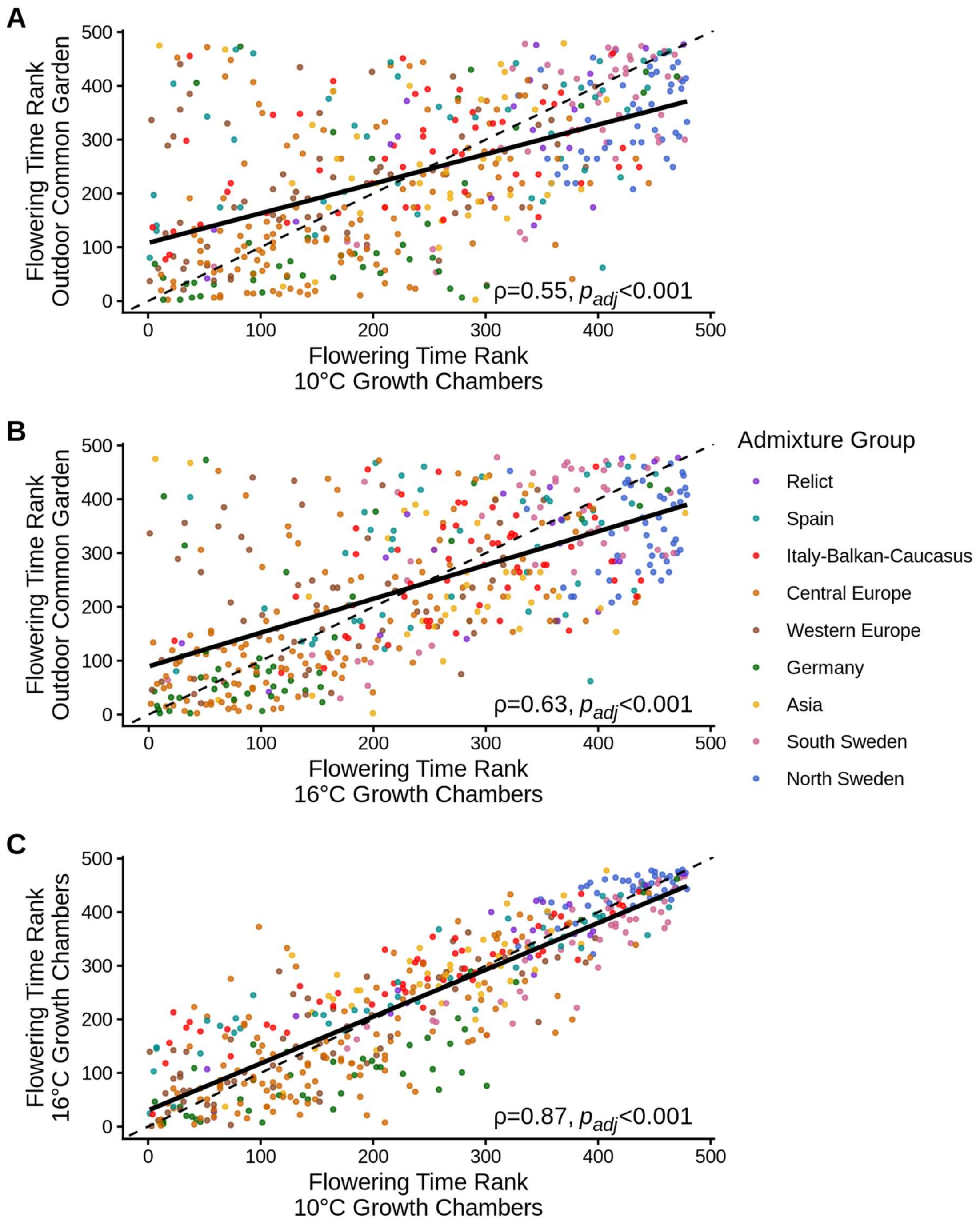


**Figure S2.** Scatter plot of the ranked mean flowering time between **A)** the outdoor common garden and 10°C growth chamber, **B)** the outdoor common garden and 16°C growth chamber, and **C)** the 10°C and 16°C growth chambers. Each data point represents a single accession colored by admixture group as assigned by the 1001 Genomes Consortium (2016). The line of best fit is the solid black line. The 1:1 line is the dashed black line. The Spearman's rank correlation and p-values adjusted for multiple comparison testing are in the bottom corner of each plot.

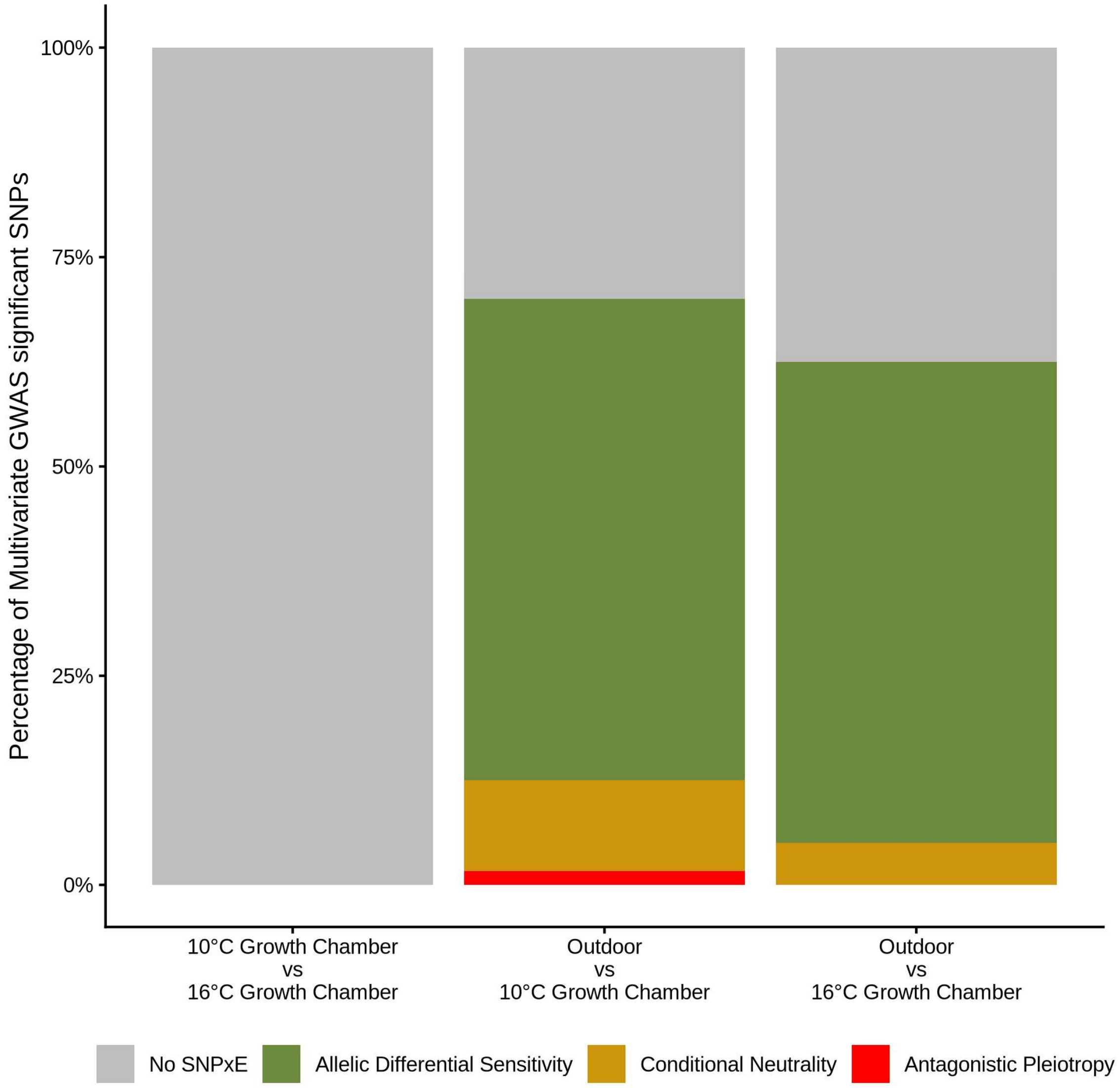


**Figure S3.** Stacked bar plot showing the percentage of significant SNPs from the multivariate GWAS that have no evidence of SNPxE, allelic differential sensitivity, conditional neutrality, and antagonistic pleiotropy between growth conditions.

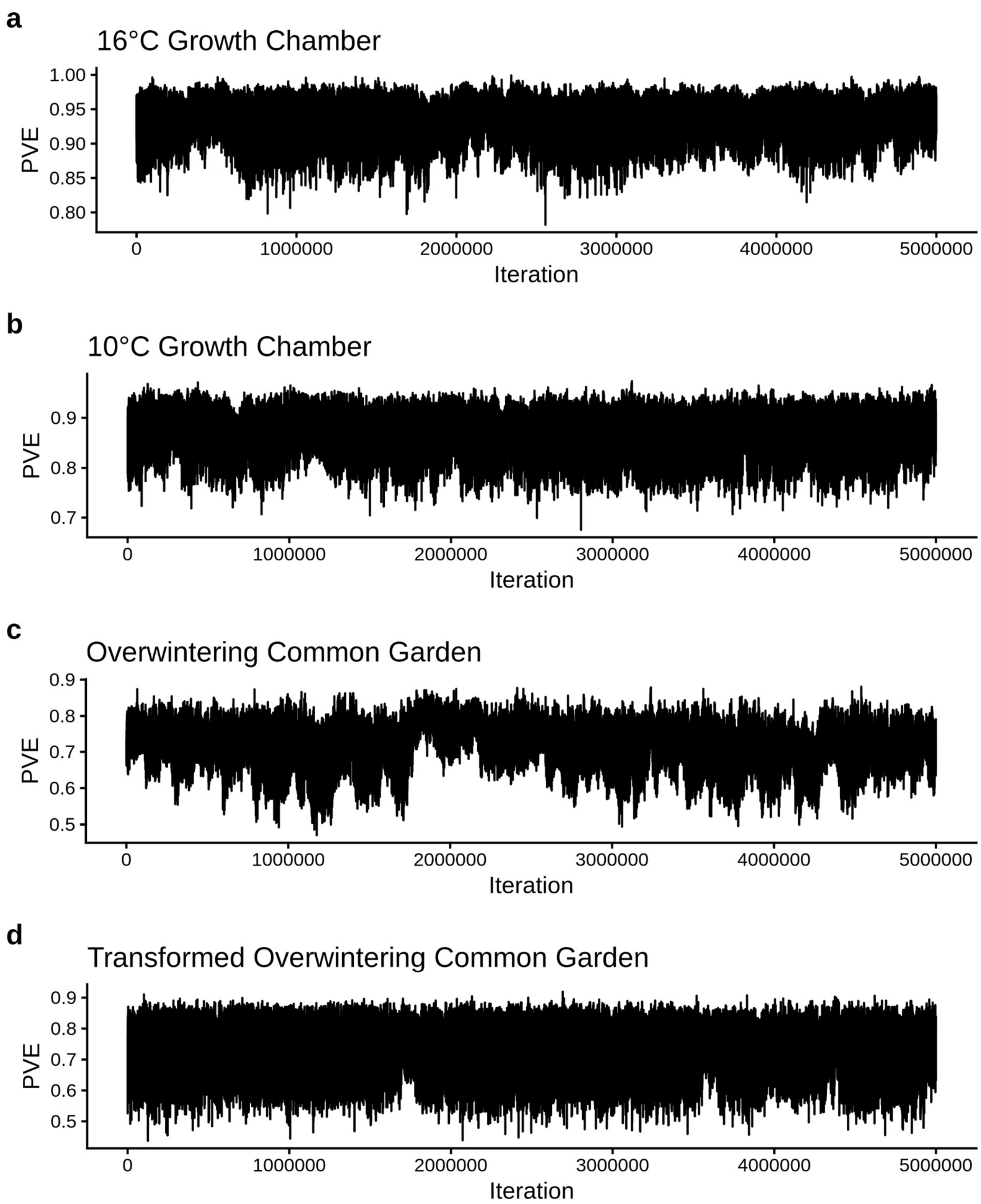


**Figure S4.** Trace plots for every 10th iteration of the BSLMM posterior distribution for proportion of phenotypic variation explained by genotype (PVE) for flowering time in the **a)** 16°C growth chamber **b)** 10°C growth chamber, **c)** untransformed outdoor common garden, and **d)** transformed outdoor common garden.

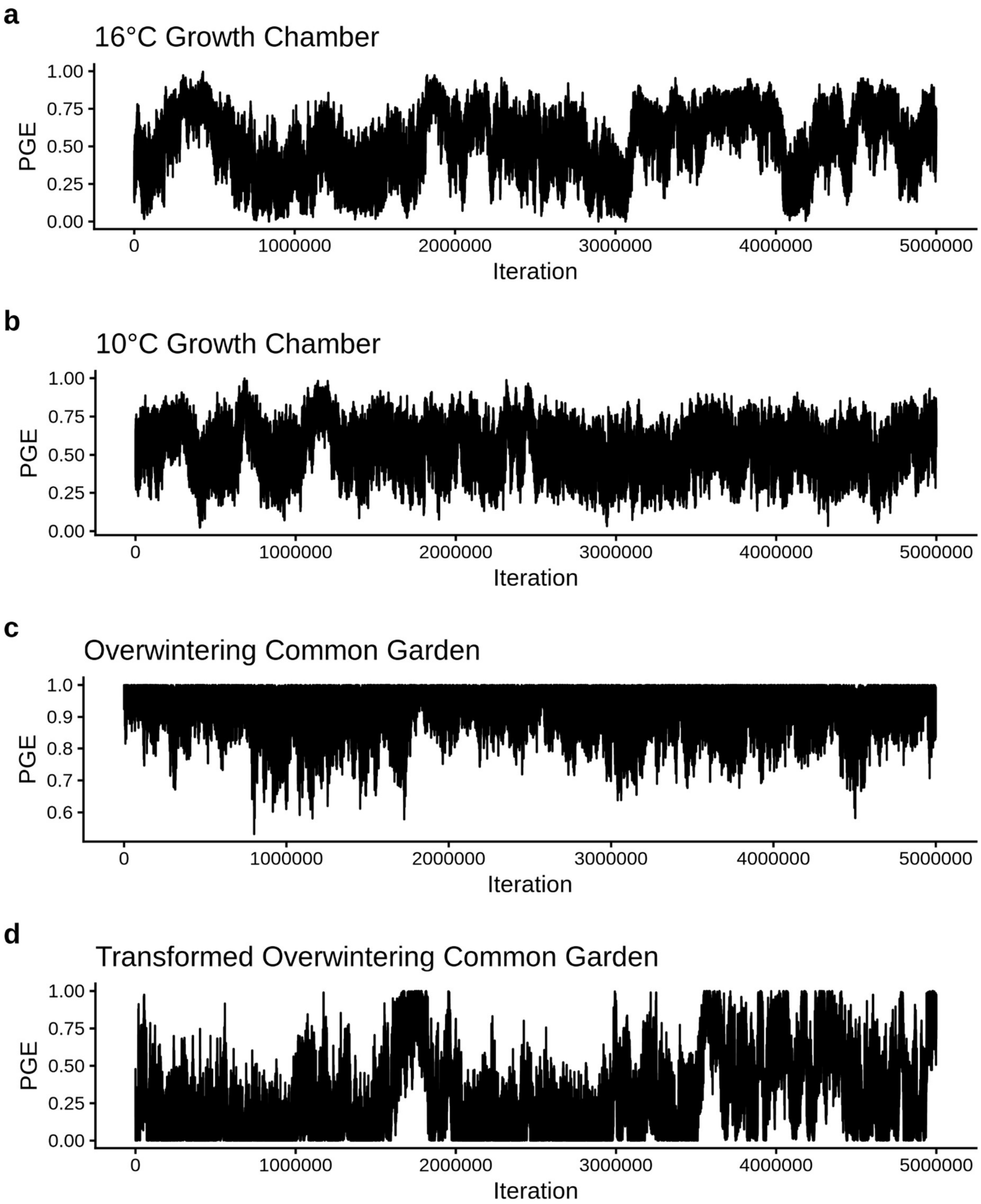


**Figure S5.** Trace plots for every 10th iteration of the BSLMM posterior distribution for proportion of genetic variance explained by large effect loci (PGE) for flowering time in the **a)** 16°C growth chamber **b)** 10°C growth chamber, **c)** untransformed outdoor common garden, and **d)** transformed outdoor common garden.

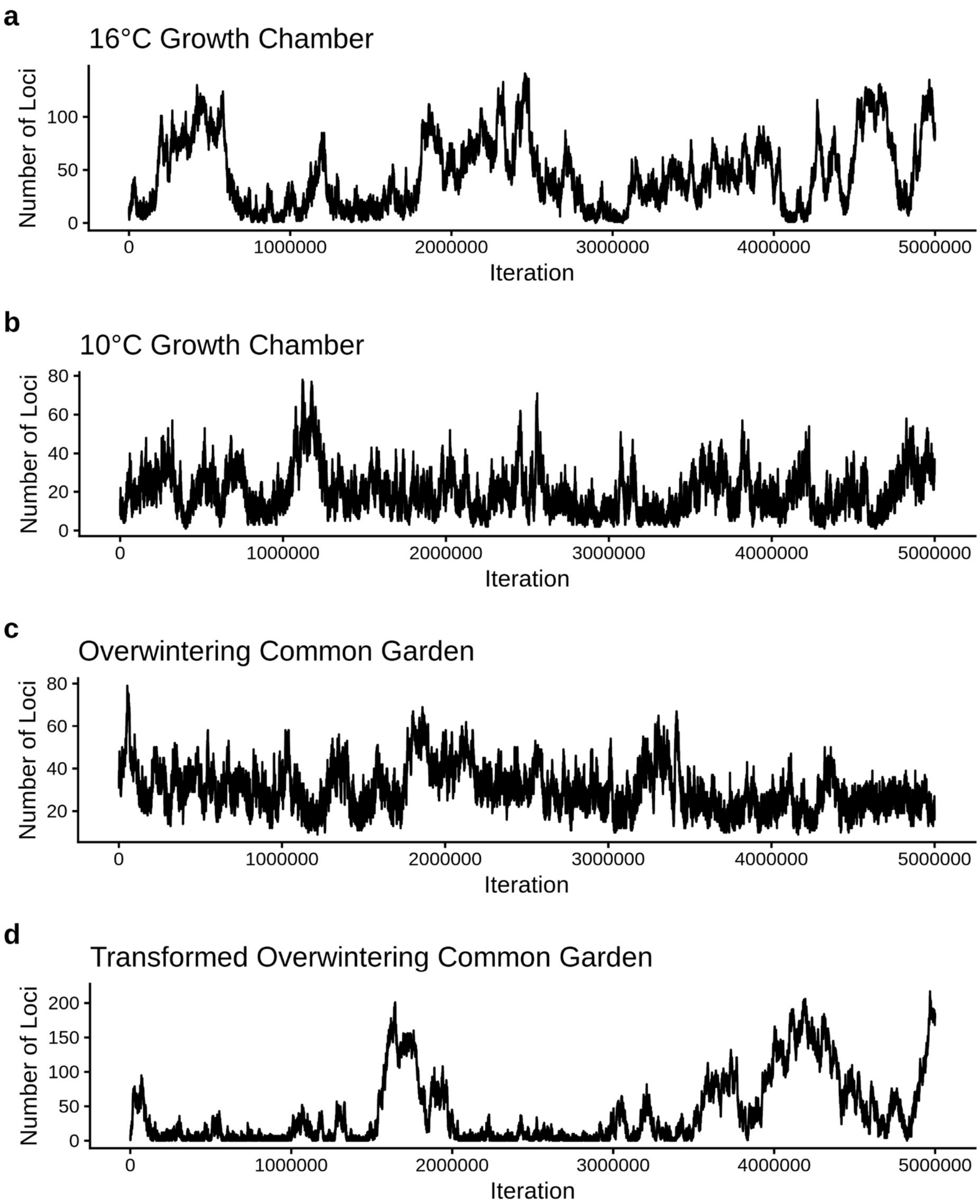


**Figure S6.** Trace plots for every 10th iteration of the BSLMM posterior distribution for number of large effect loci for flowering time in the **a)** 16°C growth chamber **b)** 10°C growth chamber, **c)** untransformed outdoor common garden, and **d)** transformed outdoor common garden.

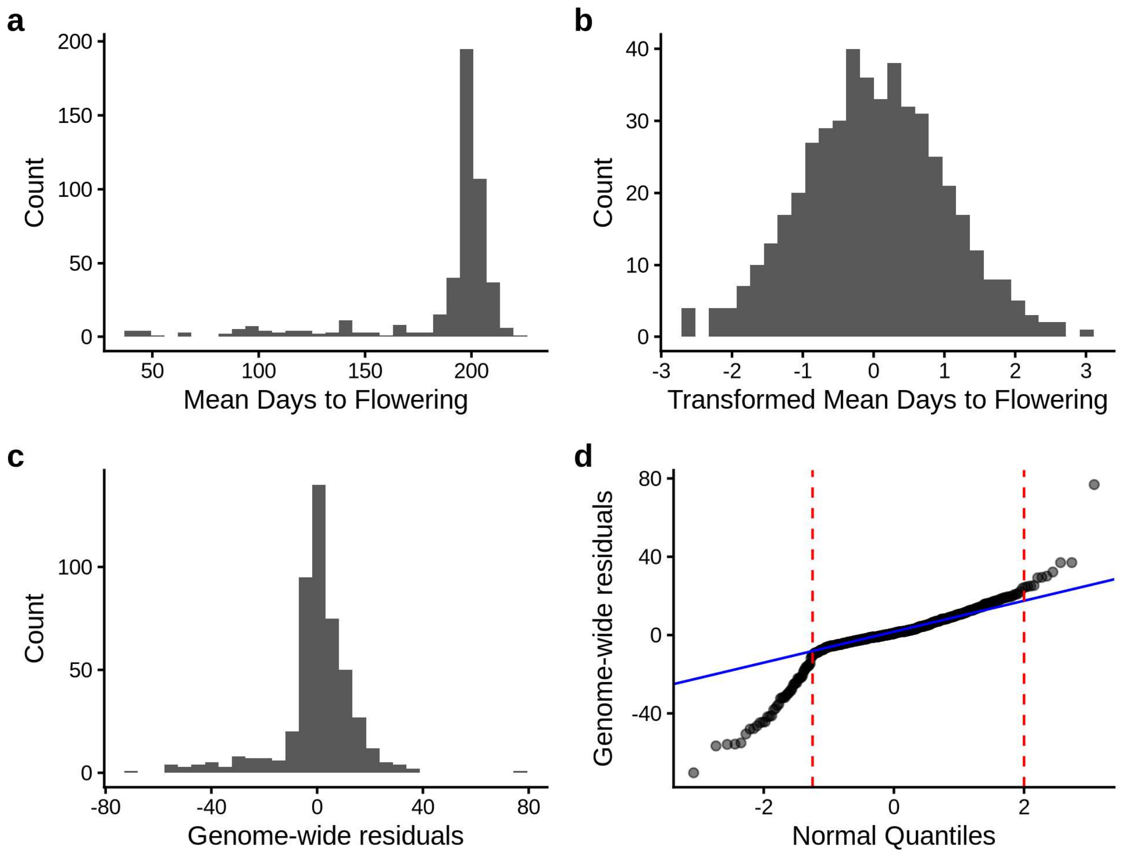


**Figure S7.** Distributions of **a)** untransformed flowering time data in the outdoor common garden, **b)** rank-based inverse normal transformed flowering time in the outdoor common gatden, and **c)** genome-wide residuals for flowering time outdoor common garden GWAS. **d)** QQ-plot for genome-wide residuals. The blue solid line is the QQ reference line that goes through the first and third quartile. Individuals within the red-dashed lines (n = 417) were used to test if skewed phenotypic distributions affect GWAS results.

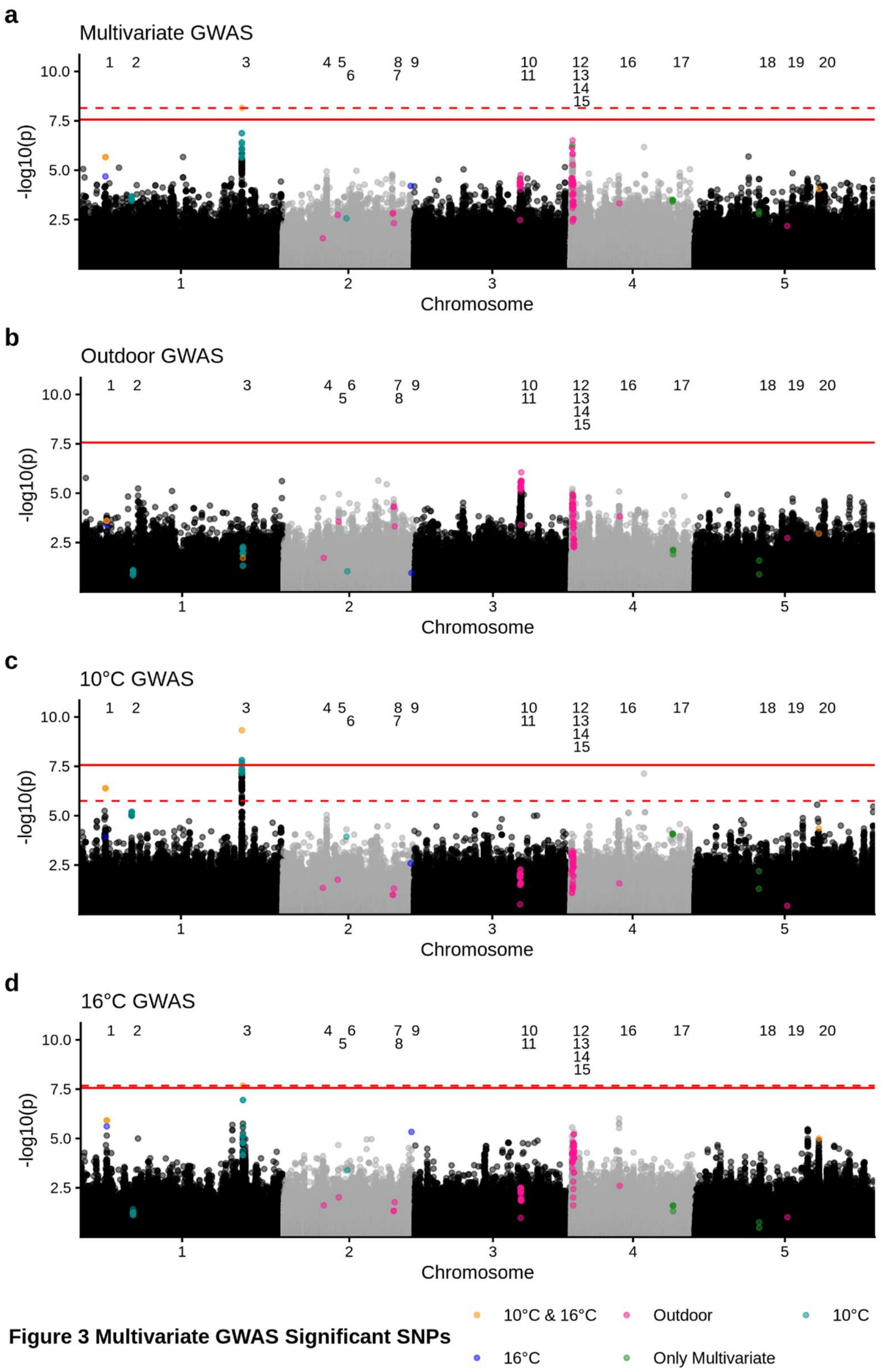


**Figure S8.** Manhattan plots for flowering time rank-based inverse normal transformed data for the **a)** multivariate GWAS, **b)** univariate outdoor common garden GWAS, **c)** univariate 10°C growth chamber GWAS, and **d)** univariate 16°C growth chamber GWAS. The red dashed line represents the largest p-value that surpasses the FDR 0.05 significance threshold (if applicable). The red solid line is the Bonferroni significance threshold. Significant SNPs are grouped into peaks labeled at the top of each Manhattan plot based with the same numerical labels as Fig 3.

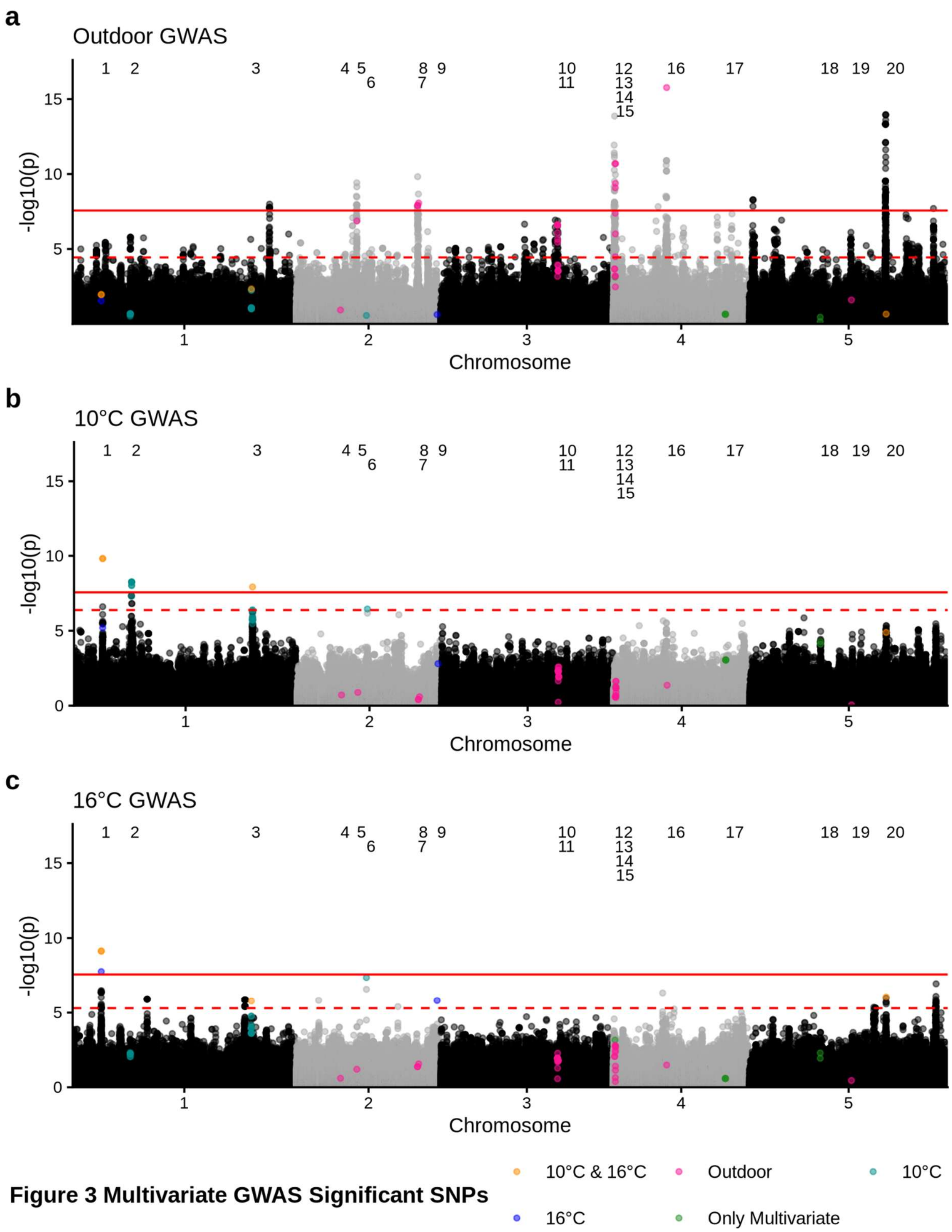


**Figure S9.** Manhattan plots for flowering time that only includes the individuals within the red-dashed lines from the qq plot in Fig S7d (n=417). **a)** univariate outdoor common garden GWAS, **b)** univariate 10°C growth chamber GWAS, and **c)** univariate 16°C growth chamber GWAS. The red dashed line represents the largest p-value that surpasses the FDR 0.05 significance threshold. The red solid line is the Bonferroni significance threshold. Locations of peaks from the multivariate GWAS in Fig 3 are labeled at the top of each Mahattan plot.

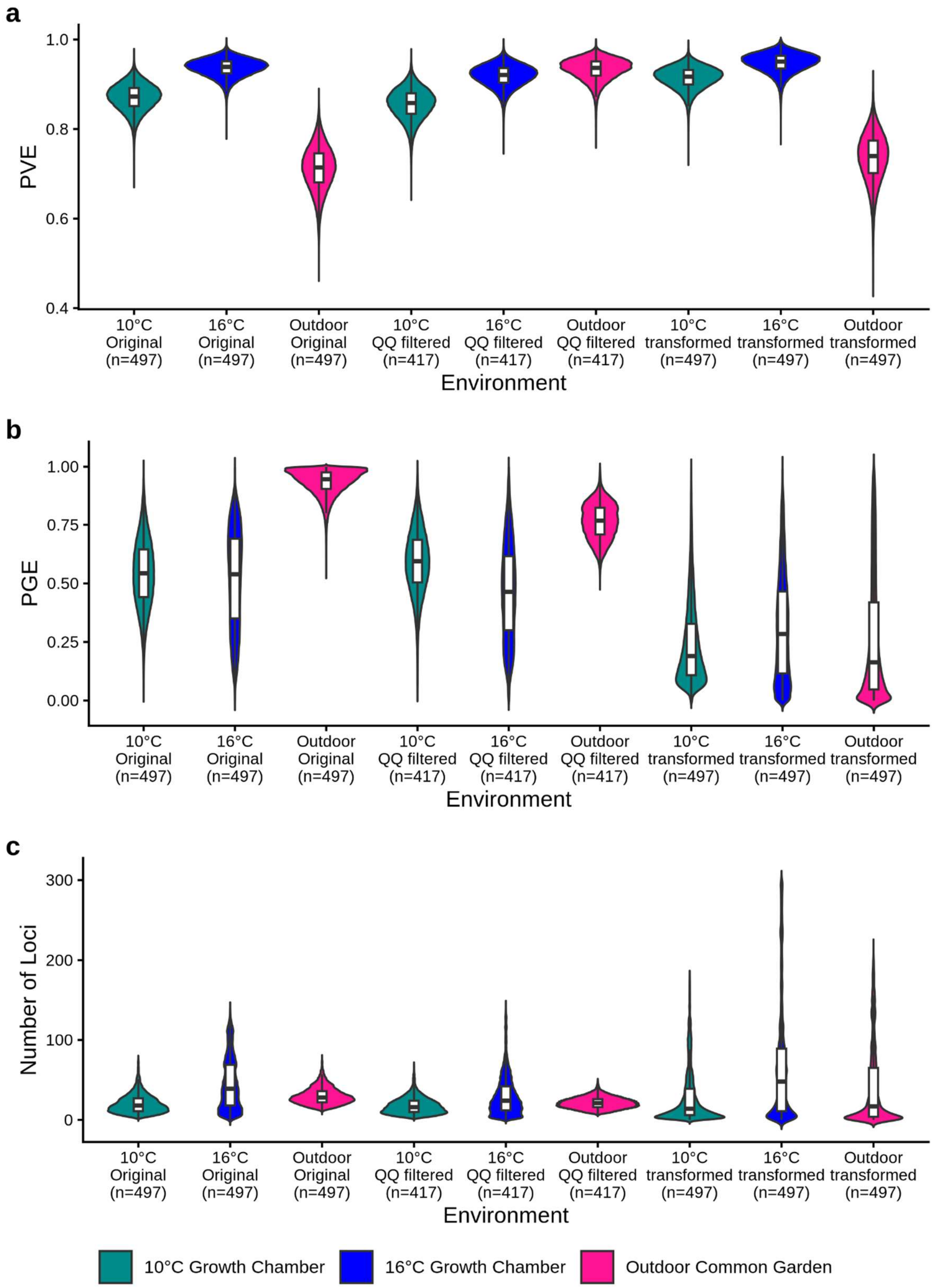


**Figure S10.** BSLMM posterior distributions for **a)** proportion of phenotype variance explained by genotype (PVE), **b)** proportion of genetic variance explained by large effect loci, and **c)** estimated number of large effect loci for the untransformed full flowering time dataset (as shown in Fig 5), the filtered dataset based on genome wide residuals (QQ filtered), and the rank-based inverse normal transformed outdoor common garden data. The violin plot color indicates the growth condition.

### Tables

Table S1: Rank based correlation (ρ) and p-values adjusted for multiple pairwise comparisons for each admixture group across growth environments. Correlations that are significant are bolded.

| **Admixture Group** | **Outdoors vs 10°C growth chamber** | | | **Outdoors vs 16°C growth chamber** | | | **10°C vs 16°C growth chamber** | | |
|---|---|---|---|---|---|---|---|---|---|
| | **ρ** | **$p_{adj}$ value** | **95% CI** | **ρ** | **$p_{adj}$ value** | **95% CI** | **ρ** | **$p_{adj}$ value** | **95% CI** |
| All accessions | **0.55** | **<0.001** | **0.47, 0.62** | **0.63** | **<0.001** | **0.56, 0.69** | **0.87** | **<0.001** | **0.85, 0.90** |
| Relict | **0.56** | **0.04** | **0.05, 0.86** | **0.58** | **0.03** | **0.05, 0.86** | **0.79** | **<0.001** | **0.37, 0.95** |
| Spain | **0.42** | **0.01** | **0.08, 0.68** | **0.43** | **0.01** | **0.10, 0.67** | **0.95** | **<0.001** | **0.90, 0.98** |
| Italy-Balkan-Caucasus | **0.37** | **0.02** | **0.08, 0.61** | **0.35** | **0.03** | **0.04, 0.59** | **0.91** | **<0.001** | **0.81, 0.95** |
| Central Europe | **0.42** | **<0.001** | **0.24, 0.57** | **0.65** | **<0.001** | **0.54, 0.73** | **0.68** | **<0.001** | **0.55, 0.77** |
| Western Europe | **0.39** | **0.01** | **0.10, 0.61** | 0.27 | 0.10 | -0.005, 0.51 | **0.81** | **<0.001** | **0.69, 0.89** |
| Germany | **0.51** | **0.001** | **0.16, 0.75** | **0.51** | **0.001** | **0.18, 0.75** | **0.81** | **<0.001** | **0.63, 0.91** |
| Asia | 0.07 | 1 | -0.30, 0.42 | 0.12 | 1 | -0.27, 0.47 | **0.70** | **<0.001** | **0.42, 0.86** |
| South Sweden | **0.56** | **<0.001** | **0.27, 0.77** | **0.44** | **0.006** | **0.11, 0.68** | **0.84** | **<0.001** | **0.67, 0.92** |
| North Sweden | **0.38** | **0.03** | **0.07, 0.62** | 0.33 | 0.09 | -0.01, 0.60 | **0.65** | **<0.001** | **0.42, 0.79** |

Table S2: Pearson's (R) and Spearman's rank (ρ) correlations of effect sizes for the top 1% of significant SNPs from the multivariate GWAS and 1% of randomly sampled SNPs. Effect sizes used for the comparison are taken from the univariate GWAS run for each individual environment. All adjusted p-values were significant ($p_{adj} < 0.001$).

| **Comparison** | **Multivariate** | | **Random** | |
|---|---|---|---|---|
| | **R** | ρ | **R** | ρ |
| Outdoors vs 10°C growth chamber | 0.38 | 0.39 | 0.27 | 0.25 |
| Outdoors vs 16°C growth chamber | 0.49 | 0.50 | 0.31 | 0.29 |
| 10°C vs 16°C growth chambers | 0.80 | 0.81 | 0.77 | 0.75 |

Table S3: Pairwise comparisons for effect sizes for multivariate GWAS peaks across environments. GWAS peaks are labeled 1-20 as denoted in Fig 3. We categorized the type of SNPxE for each peak as allelic differential sensitivity (ADS), conditional neutrality (CN), antagonistic pleiotropy (AP), or having no SNPxE. If a peak included SNPs that differed in the type of SNPxE, we listed all categories and then number of SNPs for each type in brackets.

| **Peak** | **Significant GWAS** | **Outdoors vs 10°C growth chamber** | **Outdoors vs 16°C growth chamber** | **10°C vs 16°C growth chamber** |
|---|---|---|---|---|
| 1 | 10°C and 16°C | No SNPxE | No SNPxE | No SNPxE |
| 2 | 10°C | No SNPxE | No SNPxE | No SNPxE |
| 3 | 10°C and 16°C | No SNPxE | No SNPxE | No SNPxE |
| 4 | Outdoor | CN | CN | No SNPxE |
| 5 | Outdoor | ADS | ADS | No SNPxE |
| 6 | 10°C | No SNPxE | No SNPxE | No SNPxE |
| 7 | Outdoor | CN | ADS | No SNPxE |
| 8 | Outdoor | CN | ADS | No SNPxE |
| 9 | 16°C | No SNPxE | No SNPxE | No SNPxE |
| 10 | Outdoor | ADS (18 SNPs); CN (2 SNPs) | ADS (19 SNPs); CN (1 SNP) | No SNPxE |
| 11 | Outdoor | ADS | ADS | No SNPxE |
| 12 | Outdoor | ADS | ADS (7 SNPs); No SNPxE (5 SNPs) | No SNPxE |
| 13 | Outdoor | CN | ADS | No SNPxE |
| 14 | Multiviarte only | ADS | No SNPxE | No SNPxE |
| 15 | Outdoor | ADS (30 SNPs); CN (4 SNPs) | ADS (30 SNPs); CN (1 SNP); No SNPxE (3 SNPs) | No SNPxE |
| 16 | Outdoor | ADS | ADS | No SNPxE |
| 17 | Multivariate only | No SNPxE | No SNPxE | No SNPxE |
| 18 | Multivariate only | AP | CN | No SNPxE |
| 19 | Outdoor | CN | CN | No SNPxE |
| 20 | 10°C and 16°C | No SNPxE | No SNPxE | No SNPxE |

Table S4: Shapiro Wilk normality test statistics for transformed flowering time data for the outdoor common garden. Significant p-values, which indicates that data is not normally distributed, are bolded.

| Transformation | Test statistic (W) | p-value |
|---|---|---|
| Not-transformed | **0.596** | **<0.001** |
| Log | **0.491** | **<0.001** |
| Reflected log | **0.833** | **<0.001** |
| Square Root | **0.548** | **<0.001** |
| Boxcox | **0.864** | **<0.001** |
| Rank-based inverse normal transformation | 0.999 | 0.99 |
